# The *Bacillus* phage SPβ and its relatives: A temperate phage model system reveals new strains, species, prophage integration loci, conserved proteins and lysogeny management components

**DOI:** 10.1101/2021.11.22.469490

**Authors:** Katharina Kohm, Valentina A. Floccari, Veronika T. Lutz, Birthe Nordmann, Carolin Mittelstädt, Anja Poehlein, Anna Dragoš, Fabian M. Commichau, Robert Hertel

## Abstract

The *Bacillus* phage SPβ has been known for about 50 years, but only a few strains are avalible. We isolated four new wild type strains of the *SPbeta* species. Phage vB_BsuS-Goe14 introduces its prophage into the *spoVK* locus, previously not observed to be used by SPβ-like phages. We could also reveal the SPβ-like phage genome replication strategy, the genome packaging mode, and the phage genome opening point. We extracted 55 SPβ-like prophages from public *Bacillus* genomes, thereby discovering three more integration loci and one additional type of integrase. The identified prophages resembled four new species clusters and three species orphans in the genus *Spbetavirus*. The determined core proteome of all SPβ-like prophages consists of 38 proteins. The integration cassette proved to be not conserved even though present in all strains. It consists of distinct integrases. Analysis of SPβ transcriptomes revealed three conserved genes, *yopQ*, *yopR*, and *yokI*, to be transcribed from a dormant prophage. While *yopQ* and *yokI* could be deleted from the prophage without activating the prophage, damaging of *yopR* led to a clear-plaque phenotype. Under the applied laboratory conditions, the *yokI* mutant showed an elevated virion release implying the YokI protein being a component of the arbitrium system.

## Introduction

Phages or bacteriophages are viruses of bacteria and the most abundant biological entities on our planet. After infection, they take over the host metabolism and use it to reproduce. Direct reproduction is called lytic cycle, and the respective phages are called lytic phages. Temperate phages can integrate their genetic material into the bacterial genome, inactivate their lytic gene sets, and replicate with their host as a unit. This process results in a prophage and a lysogenic bacterium. A prophage can impart new features to its host through the additional genetic material and, in rare cases, even turn it into a pathogen (Kohm and Hertel, 2021).

*Bacillus subtilis* is a Gram-positive, rod-shaped, aerobe, spore-forming bacterium mainly found in soil (Earl *et al*., 2008). The tryptophan auxotrophic strain *B. subtilis* 168 is capable of genetic competence (Spizizen, 1958), making it a model organism for many aspects of bacterial molecular biology (Sonenshein *et al*., 2001). The genome of the model strains *B. subtilis* 168 was first sequenced in 1997 (Kunst *et al*., 1997), re-sequenced in 2009 (Barbe *et al*., 2009), and faces frequent annotation updates (Belda *et al*., 2013; Borriss *et al*., 2018), making it one of the best-characterised bacterial genomes. The genomic investigations revealed one integrative and conjugative element (ICEBs1) (Auchtung *et al*., 2016), four prophage-like regions, and two prophages, known as PBSX (Seaman *et al*., 1964) and SPβ (Brodetsky and Romig, 1965). All these alien genomic elements are non-essential for *B. subtilis* 168 and can be deleted from its genome (Westers *et al*., 2003). *B. subtilis* is a key species of a species complex known as the *B. subtilis* clade or just Subtilis-clade (Fritze, 2004; Rooney *et al*., 2009). It consists of closely related species like *B. subtilis*, *B. velezensis, B. amyloliquefaciens, B. licheniformis*, *B. glycinifermentas, B. vallismortis, B. atrophaeus, B. safensis, B. sonerensis, B. pumilus* (Fan *et al*., 2017). They are mainly mesophiles and neutrophils and morphologically similar (Fritze, 2004). Members of this clade provide suitable hosts for phages initially isolated on *B. subtilis*, like φ29 (Reilly and Spizizen, 1965; Meijer *et al*., 2001), SP-15 (Taylor and Thorne, 1963) or SPO1 (Klumpp *et al*., 2010), and contain diverse SPβ-related prophages as recently demonstrated (Dragoš *et al*., 2021).

Phage SPβ was independently described twice as one of two "defective" prophages of *B. subtilis* 168 (Seaman *et al*., 1964; Hemphill and Whiteley, 1975). It resembles the *Siphoviridae* morphotype with an icosahedral head (82 to 88 nm) and a 12 nm with and 320 nm long flexible non-contractile tail, with a 36 nm wide baseplate exhibiting six equidistant, radial projections (Hemphill and Whiteley, 1975; Warner *et al*., 1977). Its prophage is 134 kb in size and integrates into the *spsM* gene. The genome of SPβ is structured in clusters I and II containing the early phage genes and cluster III the late genes (Lazarevic *et al*., 1999). With the discovery of *B. subtilis* CU1050, SPβ proved to be capable of lytic replication and lysogenisation (Warner *et al*., 1977; Johnson and Grossman, 2016). Nevertheless, SPβ was discovered and described as a prophage of *B. subtilis* 168, which itself is the result of a mutagenesis experiment to gain auxotrophic mutants (Burkholder and Giles, 1947). Therefore, it is unclear if SPβ represents a wild type phage or a mutant as part of the laboratory strain *B. subtilis* 168. Many SPβ-related phage isolates are reported in the literature but were lost over the years, making SPβ and φ3T (Tucker, 1969) the last available historical isolates. Taxonomically the ICTV (International Committee on Taxonomy of Viruses) classifies the species in the genus *Spbetavirus*, which belongs to the *Siphoviridae* family (Virus Taxonomy Release 2020: https://talk.ictvonline.org/taxonomy/).

Despite over 50 years of research on SPβ, only individual aspects of its biology are understood. The "arbitrium"-system, responsible for the lysis-lysogeny decision in SPβ-related phages (Erez *et al*., 2017; Gallego Del Sol *et al*., 2019), and its phage-host recombination system, responsible for prophage insertion into and excision out of the hosts’ chromosome (Nicolas *et al*., 2012; Abe *et al*., 2014, 2017, 2020), are two well-investigated exceptions. Aspects of the SPβ biology between prophage establishment and lytic replication, like lysogeny management and resolvement, or between prophage excision and particle release, are entirely dark matter and require scientific attention (Kohm and Hertel, 2021).

As temperate phages significantly impact the properties of their hosts and thereby meaningfully impact their ecology and evolution, it is of utmost importance to comprehensive explore the biology of this phage group. As a prophage of the model organism *B. subtilis* 168, SPβ is an excellent model system. With this in mind, we aimed to use the possibilities of the genomic era to explore the genomic diversity of SPβ-related phages, extract the core components defining this phage group, and explore their potential functions if appropriate. We isolated and sequenced new wild type SPβ phages and revealed the mode of viral genome replication and packaging employed by SPβ. Through the investigation of SPβ prophages, we found new prophage insertion locations and discovered new SPβ-related viral species. The discovery of the SPβ core genes in combination with prophage transcriptomes and deletion experiments revealed new SPβ lysogeny management components.

## Material and Methods

### Phage isolation and generation of lysogenic strains

*Bacillus subtilis* Δ6 (Westers *et al*., 2003) and *B. subtilis* TS01 (Schilling *et al*., 2018) served as host strains. Wild type SPβ-like phages were isolated using the agar overlay plaque technique as described previously (Willms *et al*., 2017). Agarose overlay consisted of LB medium supplemented with 0.4 % (w/v) agarose. Sterile filtered raw sewage water from the Göttingen municipal sewage plant (Göttingen, Germany, 51°33’15.4" N 9°55’06.4" E) served as a source for phage isolation. All four isolated phages were submitted to the public "German Collection of Microorganisms and Cell Cultures GmbH" (DSMZ) and thereby made avalible to the scientific community.

Lysogens were isolated from turbid plaques by picking those with a sterile toothpick and resuspending the host cells in sterile LB (10 g * L^-1^ tryptone; 5 g * L^-1^ yeast extract; 10 g *L^-1^ NaCl (Miller, 1972)). Dilutions of this suspension were spread on LB agar plate. Single colony forming units (CFU) were inoculated in 4 ml LB medium and cultured for ∼ 16 h at 37 °C at vigorous shaking. Cells were removed by centrifugation 13,000 g*1 min^-1^. The supernatant was used for agar overlay plaque assay with *B. subtilis* Δ6 or *B. subtilis* TS01 to verify the spontaneous release of viral particles from the present prophage. Observed plaques confirmed the presence of a lysogen.

### Phage genome sequencing

Phages were sequenced from phage DNA and as prophages from chromosomal DNA of their lysogens. In the case of direct phage DNA sequencing, phages were first singularized via an overlay plaque assay. A single plaque was picked with a sterile toothpick and resuspended in sterile 500 µl LB. The obtained phage suspension was used to infect a 4 ml LB culture of a susceptible host at the logarithmic growth with an OD_600_ of ∼ 0.8 and incubated at 37 °C at vigorous shaking until total lysis of the culture. The lysed culture was centrifugated at 5,000 g for 5 min to pellet remaining cells and cell debris. The phages in the supernatant were sterile filtered with an 0.45 µl syringe filter (Sarstedt, Nümbrecht, Germany), supplied with 5 U/ml salt active nuclease (SERVA Electrophoresis GmbH, Heidelberg, Germany) and incubated for ∼ 16 h at 8 °C to remove free nucleic acids. At the same incubation period, the phages were also precipitated by setting the suspension to 0.5 M NaCl and 10 % (w/v) polyethylene glycol (PEG) 6000 (Sigma-Aldrich, Taufkirchen, Germany). Precipitated phages were pelleted with 14,000 g for 30 min, the supernatant discarded, and the phage pellet used for phage DNA preparation with the MasterPure complete DNA and RNA purification kit (Epicentre, Madison, WI, USA).

Lysogenic bacteria were grown in a 4 ml LB medium and cultured for ∼ 16 h at 37 °C at vigorous shaking, 1 ml of this culture was pelleted, and the cells were used for chromosomal DNA preparation. Phage and bacterial genomic DNA were prepared with the MasterPure complete DNA and RNA purification kit (Epicentre, Madison, WI, USA).

Illumina paired-end shotgun libraries were prepared with the NEBNext Ultra II FS DNA Library Prep (New England Biolabs GmbH, Frankfurt, Germany) for SPβ, Goe11 and Goe14 and Nextera XT DNA Sample Preparation Kit (Illumina, San Diego, CA, USA) for Goe12, Goe13 and lysogens and sequenced with the MiSeq system and reagent kit V.3 (2 x 300 bp) (Illumina, San Diego, CA, USA) and the NovaSeq system (2x 150bp) with GENEWIZ, Leipzig, Germany.

Raw reads were quality analysed with FastQC (https://www.bioinformatics.babraham.ac.uk/projects/fastqc/) and, if necessary, quality processed with Trimmomatic 0.39 (Bolger *et al*., 2014). All obtained sequences were submitted to the SRA archive. Respective accession numbers can be found in supplementary materials 1.

### Genome assembly and annotation

Phage genomes were assembled with the Unicycler v0.4.8 pipeline (Wick *et al*., 2017) employing SPAdes version: 3.14.0 (Bankevich *et al*., 2012) and accepted as complete when resulting in a circular replicon. The final phage genome was orientated like the SPβ c2 genome (Lazarevic *et al*., 1999). The protein-coding genes were initially predicted and annotated with the Prokka 1.14.5 pipeline (Seemann, 2014), complemented with an InterProScan5 (Jones *et al*., 2014) protein domain search and finally manually curated. Individual proteins were also investigated with the web-based InterProScan version (https://www.ebi.ac.uk/interpro/).

The genome of the Goe14 lysogen was artificially constructed *in silico*. Therefore, the viral sequence reads were mapped to the genome sequence of *B. subtilis* Δ6 [NZ_CP015975] using the bowtie2 read aligner version 2.2.6 (Langmead and Salzberg, 2012) with the option –local. The created alignment was visualised with Tablet 1.17.08.17 (Milne *et al*., 2010), and so the hybrid read, consisting of host and virus sequence, was identified (supplementary materials 2). The genome of the lysogen was created by integrating the viral genome of Goe14 into the genome of *B. subtilis* Δ6 [NZ_CP015975]. The resulting prophage integration was consistent with the prophage of *B. subtilis* BS155.

### Phage packaging mechanism and packaging start point determination

Phage packaging mechanism and packaging start points were determined with the PhageTerm software package running on a Galaxy instance of the Pasteur Institute (https://galaxy.pasteur.fr). Raw Illumina reads, generated with a NEBNext Ultra II FS DNA Library Prep (New England Biolabs GmbH, Frankfurt, Germany) sequence library, were used as input.

Identification of inverted repeats in the genome fragment containing the SPβ pac-site was realized with the RNAfold Web Server (http://rna.tbi.univie.ac.at/cgi-bin/RNAWebSuite/RNAfold.cgi) using standard parameters and the DNA parameters (Gruber *et al*., 2008). The multiple sequence alignment was realized with the Clustal Omega algorithm made available on the EMBL-EBI website (https://www.ebi.ac.uk/Tools/msa/clustalo/) (supplementary materials 11)

### Extraction of SPβ-like prophages

Draft genomes of *B. subtilis* strain CU1065(Z) [NZ_JADOXU010000001], and *B. subtilis* strain DBS-15(rho11) [JADOXP010000001] were downloaded from GenBank and alight to the genome of *B. subtilis* 168 [NC_000964] with Mauve 20150226 build 10 for Windows (Darling, 2004). Contigs containing the SPβ-related prophage were identified and extracted using Artemis Release 18.1.0 (Carver *et al*., 2011). The so obtained prophage genomes of Z and ρ11 were compared to SPβ using Mauve 20150226 build 10 for Windows (Darling, 2004).

The bioinformatic identified SPβ-related prophages by Dragoš and co-workers (Dragoš *et al*., 2021) served as a starting point for curating and extracting new SPβ-related prophages. Derivate strains of *B. subtilis* 168 were excluded. Prophages that were known to reside in *spsM* and *kamA* were extracted from the host-genomes using Artemis Release 18.1.0 (Carver *et al*., 2011) and, if appropriate, orientated like SPβ c2 [NC_001884] (Lazarevic *et al*., 1999) for comparative analysis.

To identify new integration loci, genomes of the lysogens were directly compared with the genome of *B. subtilis* 168. A BLASTn comparison file was created by using makeblastdb and BLASTn version 2.10.0 (Altschul *et al*., 1990). The comparison was visualised with ACT Release 18.1.0 (Carver *et al*., 2011). The SPβ-like prophages were tracked and identified through their homology to SPβ specific genes. The prophage boundaries were predicted through the presence of a site-specific recombinase gene. The integration locus and the exact *attP* and *attB* sites were identified through sequence comparison of gene fragments flanking the identified prophages with an intact counterpart in *B. subtilis* 168 or closely related strains. Those were identified via BLASTn against the NCBI nr database. The *attB* sites were defined when they fit the intact template, and *attP* sites were defined as a deviating variant of the *attB* site. For integration loci, where *attP* and *attB* matched each other, the instance located close to the viral integrase was considered *attP*. All this was realised with the bioinformatical tools Artemis Release 18.1.0 (Carver *et al*., 2011) and Mauve 20150226 build 10 for Windows (Darling, 2004).

### Prophage genome re-annotation

We used the prokka pipeline 1.14.5 (Seemann, 2014) with standard parameters to re-annotate all previously identified prophages. This procedure provided uniform protein annotation for all genomes and was particularly required as some host *Bacillus* genomes had no protein annotation, like *B. velezensis* W1 [CP028375].

### *Spbetavirus* core genome identification

Protein sequences from the re-annotated genomes were used as input for orthology detection with Proteinortho6 (Lechner *et al*., 2011). The calculation was performed with standard parameters employing DIAMOND v0.9.29.130 (Buchfink *et al*., 2015) for protein-protein comparison. MS Excel was used for data evaluation.

Proteins of the remnant prophage from *B. subtilis* 168 integrated into the *glnA* locus were compared to the proteins of the identified prophage in the same manner.

### Transcriptome analysis

The microarray-based transcriptome data for SPβ were directly extracted from Table S2 of the study by Nicolas et. al. (Nicolas *et al*., 2012). The RNA-seq based SPβ transcriptome data were obtained by the following procedure. Therefore, we used sequence data from Popp et. al. (Popp *et al*., 2020) and Benda et. al. (Benda *et al*., 2021). Raw transcriptome sequence reads were downloaded from the SRA archive using fastq-dump from sra-tools (https://github.com/ncbi/sra-tools). They were reverse complemented with the fastx_reverse_complement script from the FASTX-Toolkit (http://hannonlab.cshl.edu/fastx_toolkit/commandline.html) and mapped with to the sequence subset from *B. subtilis* 168 containing the SPβ prophage. The resulting SAM file was transformed to a tds-file and analysed with the TraV program (Dietrich *et al*., 2014). TraV was also used to calculate nucleotide activities per kilobase of exon model per million mapped reads (NPKM). Those NPKM-values represent the normalised transcriptional activity for all protein-coding genes, which TraV could extract from the given GenBank file. Regardless of the origin of the transcription activity data, an average value for each transcription was formed and used as a demarcation line for present transcriptional activity at the dormant prophage. Genes exciding the average transcription value were considered as transcriptionally active.

### Molecular cloning

If not explicitly mentioned, used molecular biological methods based on Sambrook and Russell’s method collection (Sambrook and Russell, 2001). If not otherwise stated, all chemicals were dissolved in deionised water and autoclaved at 121 °C and 2 bars for 20 min. LB media was supplemented with 1.5 % (w/v) agar-agar for solid agar plates and supplemented with respective antibiotics if required. Liquid cultures were grown in 4 ml LB in glass tubes and vigorously shaken to ensure good aeration.

*Escherichia coli* DH10B was used as a cloning strain (Durfee *et al*., 2008). Chemical competent cells were prepared with the CaCl_2_ method (Sambrook and Russell, 2001). Enzymes required for cloning were obtained from ThermoFisher Scientific, Germany and used as recommended by the manufacturer.

*B. subtilis* strains were made competent as described by Anagnostopoulos and Spizizen (Anagnostopoulos and Spizizen, 1961) with modifications (Kohm *et al*., 2021). *B. subtilis* mutants were created either through the transformation of the recipient stain with chromosomal DNA, a specific plasmid or PCR product. The Phusion™ High-Fidelity DNA Polymerase (2 U/µl) and HF buffer (ThermoFisher Scientific, Germany) were used as recommended by the manufacturer for PCR amplification from bacterial or viral chromosomal DNA. PCR products were analysed with a horizontal 1 % TAE agarose gel electrophoresis system (Sambrook and Russell, 2001). Specific PCR products, like the amplifications of the *yopR* gene from the SPβ clear plaque mutants, were sequenced with Microsynth SeqLab (Göttingen, Germany). Used primers, strains and plasmids are listed in supplementary materials 3, 4, and 5.

### Sublancin assay

The sublancin assay verified the presence of SPβ in a lysogen. In brief, 3 µl of an overnight culture of the strains to be analysed was dropped onto a lawn of sublancin sensitive *B. subtilis* strain TS01 and incubated overnight at 37 °C. A zone of growth inhibition around the strain of interest indicated the presence of the SPβ prophage.

## Results

### Historical strains and new Isolates

Many SPβ-related phage isolates like IG1, IG3, IG4 (FERNANDES *et al*., 1986) φ3T (Tucker, 1969), Z (FERNANDES *et al*., 1986), ρ11 (Dean *et al*., 1976), SPR (Noyer-Weidner *et al*., 1983) and H2 (Zahler, Korman, Thomas, and Odebralski, 1987) are reported in the literature. However, only a few are sequenced and still available as an active sample. The type phage SPβ was sequenced twice as a prophage of the model organism *B. subtilis* 168 and a heat-inducible c2 mutant. The phage φ3T was sequenced during the investigation of the arbitrium system (Erez *et al*., 2017). Recently, further historical SPβ-like isolates were sequenced through the whole genome sequencing of their lysogens. In this way, the genome of phage Z [NZ_JADOXU010000001], ρ11 [JADOXP010000001], and H2 [CP041693] were made available (Table 1).

**Table 1:**
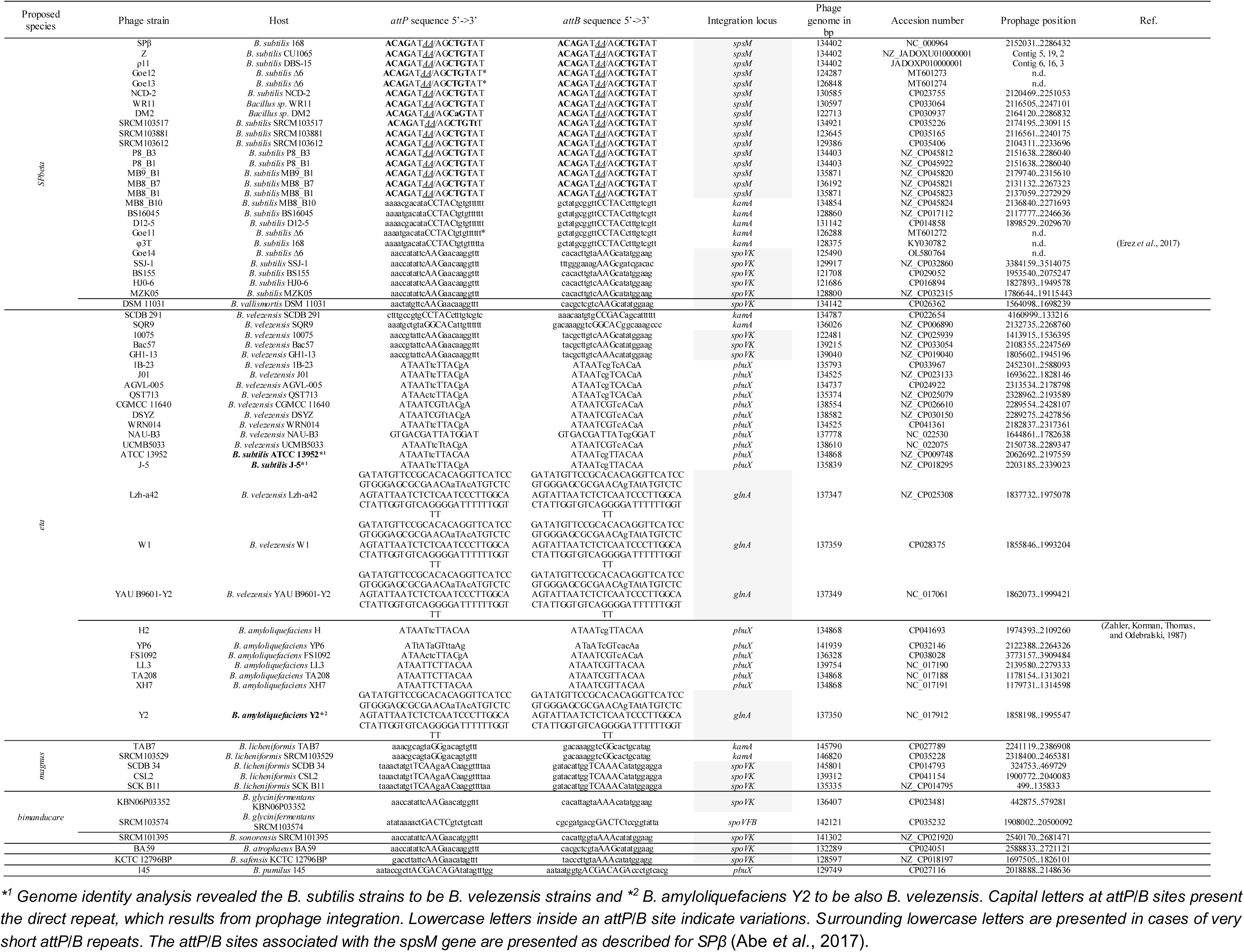
New SPβ-like phages

Besides the historical phages, we isolated four new SPβ-like environmental strains named vB_BsuS_Goe11 (Goe11), vB_BsuS_Goe12 (Goe12), vB_BsuS_Goe13 (Goe13), and vB_BsuS_Goe14 (Goe14). Whole-genome sequencing of those phages revealed Goe12 representing the smallest and Goe13 the largest genome, respectively (Table 1). BLASTn based average nucleotide identity analysis with all as functional assumed SPβ-like phage genomes revealed all phages of the same species apart of H2. Phage H2 represents a distinct species closely related to SPβ (Figure 1 A and Table S1).

**Figure 1:**
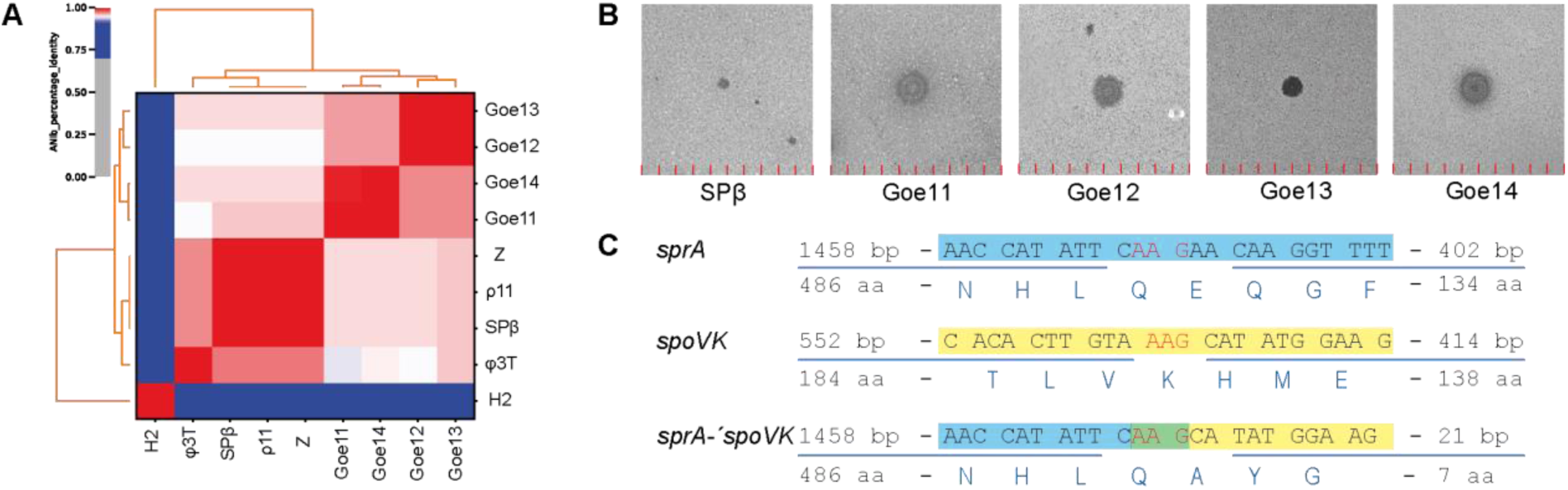
**A:** Whole-genome average nucleotide identity analysis of experimentally verified SPβ-like phages. White to red indicates identity values between 95 to 100 % and thus strains of the same species. White to blue indicates values 95 to 70 % and thus strains of the same genus. **B:** Plaques of new wild type isolates in comparison to the type strain SPβ. The plaques are presented in one square centimetre section. The red lines are a millimetre scale. **C:** Goe14 integration into the spoVK gene of B. subtilis. The sprA-like phage integrase, presented in blue, recombines the attP site (AAG) located in its coding sequence with the attB site in the spoVK gene, where those three bases present a triplet coding for lysine. Upon recombination, the sprA-like integrase gene of Goe14 alters its C-terminal coding region. It becomes significantly shorter.

The new wild type isolates formed significantly larger plaques on overlay agar, indicating more efficient reproduction than the original SPβ (Figure 1 B). Phage Goe13 revealed almost clear plaques. However, it is still can able to establish lysogeny like the remaining SPβ-like phages. All in all, the new isolates present new wild type SPβ-like phages, which can serve as model systems in the investigations of SPβ-like phage biology.

### Goe14 forms a new type of prophage

The genome sequences revealed Goe11 to contain an *attP* site similar to φ3T, and Goe12 and Goe13 similar to SPβ. Sequencing of Goe11 and Goe12 lysogens proved the integration of the prophages into *kamA* and *spsM* genomic loci, respectively (data not shown). Goe14 revealed an unknown *attP* site. However, as this strain successfully lysogenised *B. subtilis,* we assumed a potentially new integration locus for this phage. By searching the Goe14 raw reads for phage-host hybrids (supplementary materials 2), we could identify the *spoVK* gene of *B. subtilis* as a potential integration locus of Goe14. Like *kamA* and *spsM* (Feucht *et al*., 2003), *spoVK* is a sporulation associate gene (Fan *et al*., 1992; Eichenberger *et al*., 2004). Reconstruction of the Goe14 lysogen and the BLASTn based investigation of the transition area (200 bp host plus 200 bp phage) led to the identification of a 100% identical sequence pattern in the genome of *B. subtilis* BS155 [CP029052.1]. The *B. subtilis* BS155 chromosome revealed an SPβ-like prophage integrated into the *spoVK* gene, as observed with Goe14. With both data sets in hand, we recovered the *attP* and *attB* (*attP/B*) sites of Goe14 and the BS155 prophage. The duplicated direct repeat consists of the three bases AAG, which is a rather short sequence compared to the *attP/B* site of SPβ (ACAGATAAAGCTGTAT). The minimal size explains why we were not able to identify the *attP/B* site of BS155 with a prophage prediction tool like PHASTER (Arndt *et al*., 2016). However, the size is not unusual as the *attP/B* sites of φ3T also consist of only the five bases CCTAC (Suzuki *et al*., 2020). In the *spoVK* gene, the AAG triplet codes for a lysine triplet. On the Goe14 genome, the *attP* site locates in the C-terminal coding region of the phage integrase, the equivalent to *sprA* from SPβ. Upon integration, the integrase gene undergoes truncation (Figure 1 C). As no *sprB* gene equivalent was observed in the Goe14 genome, one could speculate the truncation of the integrase to fulfil a regulatory function. The extended version present on the circular phage genome could be responsible for the integration and its truncated prophage form for the excision of the prophage upon reactivation for lytic replication. In such a case, the integrase gene is likely to be transcriptionally regulated and not constitutively expressed like with SPβ (Abe *et al*., 2014).

Taken together, the genome of Goe14 reveals a third integration locus of SPβ-like phages into the chromosome of *B. subtilis* and potentially a new regulatory circuit, which controls integration and excision of the phage.

### Genome replication, opening and packaging

Previous endonuclease restriction studies of SPβ genomic DNA extracted from viral particles indicated the location of the genome opening site and packaging starting point between *yonR* (BSU_21020) and *yonP* (BSU_21030) (Fink and Zahler, 1982). Analysing raw sequence reads of four SPβ-related isolates confirmed the genome-opening sites to locate between *yonR* and *yonP.* Furthermore, we precisely identify their genome opening points (Figure 2, right panel). An alignment of the first ∼250 bp from all four genomes revealed a consistent starting point of the particle-packed genome for three out of four genomes. The genome opening point of Goe11 deviates. Looking at the coverage plots of Goe11, we can see that it does not have a sharp edge of the aligned reads, which is needed for genome opening point prediction (Figure 2 C, right panel). We take this as an indication of the automatic prediction to be slightly imprecise and the actual opening point to consistent with the three identified.

**Figure 2:**
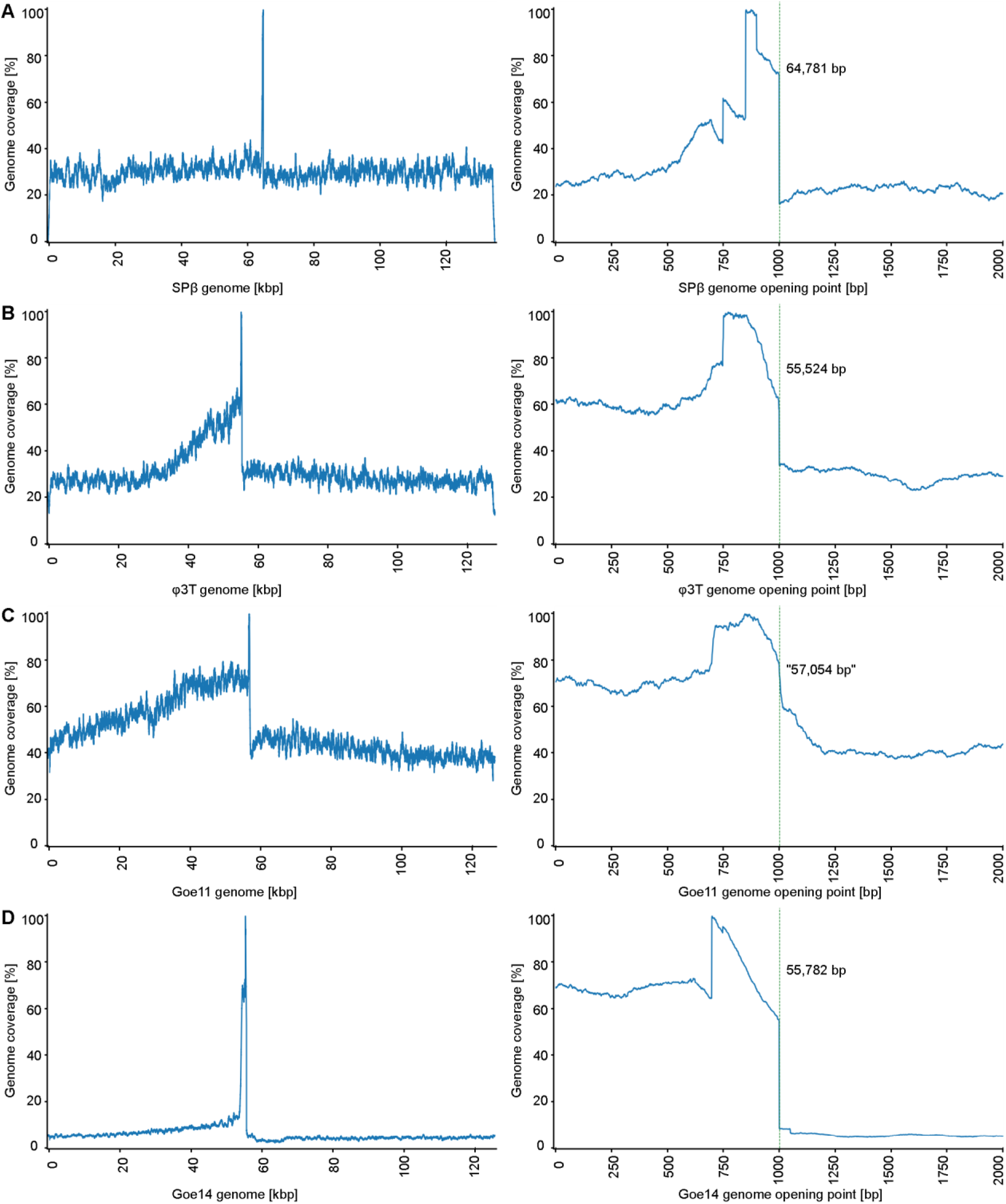
Phage genomes coverage plots. Left panels show the distribution of raw sequence read over the respective phage genome. The counter-clockwise fate out of read coverage from the coverage peak indicates a pac-site as the initiation point of genome packaging and a head-full mechanism of a concatemeric template. The packaging direction corresponds to the read fate out. Right panels present zooms into the coverage peak, which presents the respective genome opening points and the predicted first base packed into the phage head. The predicted opening point of Goe11 is set in quotation marks as we expect it to be slightly imprecise.

The raw read mapping also indicated the potential mode of genome replication and packaging employed by the SPβ-like phages. For φ3T, we can see that after the mapping peak, which represents the genome opening point, reads strongly accumulate at its left side and fade out with increasing distance from the opening point. From this observation, we can first conclude that the phage genome is packed counter-clockwise. Second, the phage genome has to be concatemeric, allowing it to fill 100 %-plus into the phage head. The observed fade out over a relatively long distance, best observed with Goe11 (Figure 2 C, left panel), indicates a head-full packaging mechanism and a circular permutation of the particle packed viral genomes. The degree of circular permutations in a viral population seems to be strain specific. No read accumulation over a significant distance was observed for SPβ, indicating almost no circular genome permutations packet into its virions. The φ3T phage reveals an evident read accumulation, and Goe11 the most pronounced. The remaining question is, what size is the "plus" which results from the head-full packaging and leads to the circular permutation. As this information is impossible to extract from the given sequence data, it has to be experimentally explored in future.

### SPβ-like prophage diversity and their *attB* sites

In times of the genomic era, many bacterial genomes are publicly available. A recent bioinformatic search for SPβ-related prophages in publicly available complete *Bacillus* genomes by Dragoš and co-workers revealed additional SPβ-related prophage candidates (Dragoš *et al*., 2021). We reviewed this list of potential candidates and extracted the prophage genomes of all lysogens manually. Thereby we considered only prophages with a genome size of more than 120 kbp that contained a *sprA-*like recombinase gene which allowed a precise determination of its boundaries. In total, 55 additional SPβ- like prophages were recovered, distributed among ten host species of the Subtilis-clade (Table 1). We recovered *pbuX*, *glnA*, *spoVFB* as new integration loci and several previously unknown *attP/B* sites (Table 1). All genes serving as integration loci for SPβ-like phages reside at the replication termini of a *Bacillus* genome (Figure 3).

**Figure 3:**
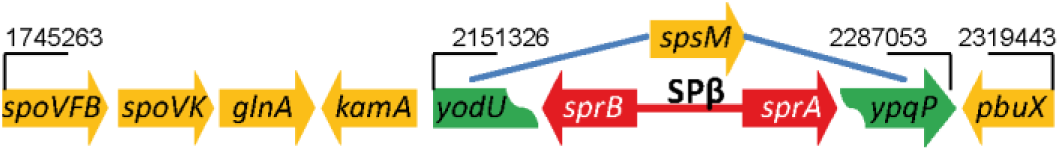
Identified integration loci for SPβ-like phage present in B. subtilis 168.

Only the *spoVFB* gene seems to be associated with sporulation (Steil *et al*., 2005). The gene *pbuX* is involved in xanthine metabolism (Christiansen *et al*., 1997), and *glnA* encodes the glutamine synthetase (Commichau and Stülke, 2008). The data also revealed that SPβ-like phages could use distinct regions of the same gene for its integration. That becomes particularly obvious with the *kamA* gene, which reveals three different *attB* sites with *B. subtilis*, *B. velezensis* and *B. licheniformis.* Multiple *attB* sites can also be observed with *spoVK* gene with *B. licheniformis* and *pbuX* in *B. pumilus* and the others. Phages integrating into *kamA* of *B. licheniformis* reveal only two guanine bases (GG) as duplication of their *attP/B* site, which is even smaller than in the case of Goe14. The *glnA* locus represents the opposite extreme. Its *attP/B* sites duplicate 106 bases upon integration (Table 1). It is also the only gene that remains intact after prophage integration. Interestingly, we observed a prophage-like element in the genome of *B. subtilis* 168 directly associated with *glnA*. Even no duplication of the 3’ end of the *glnA* gene was present, a ∼9 kbp region with a reduced CG content till *xynP* (BSU_17570) can be observed. This region contains 11 protein-coding genes, of which five showed similarities to SPβ-like proteins of the here identified prophages. The *ynzG* gene product (BSU_17490) is similar to the SunI protein of SPβ (BSU_21490) with 81 % query cover and 32.35 % identity, and YnaB (BSU_17500) revealed itself as an identical but shorter version of the SPβ YokH (BSU_21590). These results conclude that *B. subtilis* 168 hosts two SPβ-like prophages. One still functional prophage, known as SPβ and associated with the *spsM* gene and one almost degenerated and associated with the *glnA* gene.

### The genus *Spbetavirus* contains seven species

The availability of 64 SPβ-like genomes allows for the first time a holistic taxonomic analysis of these phages. The base for this investigation was an average nucleotide identity analysis employing BLASTn for pairwise comparison (Figure 4 and Table S2). Results revealed all genomes to be of the *Spbetavirus* genus, which resolves into seven species clusters.

**Figure 4:**
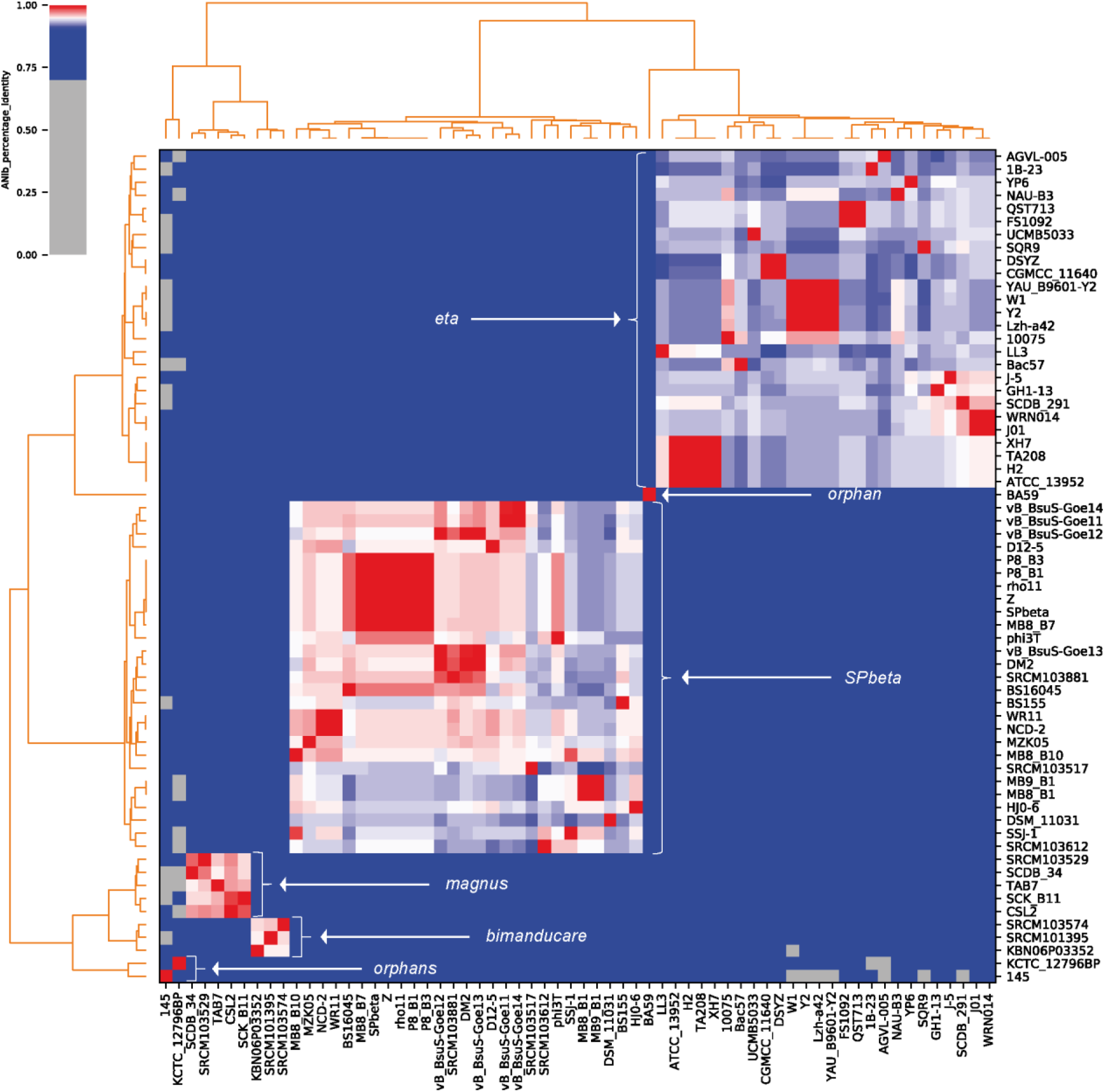
Average nucleotide identity analysis of SPβ-like phages. Analysis reveals seven viral species clusters.

The largest species cluster consists of 27 strains includes the type strain SPβ. Phage genomes of this cluster reveal an average nucleotide identity value between 95 and 100 %, typical for members of the same species (Nordmann *et al*., 2019; Baena Lozada *et al*., 2020). To be precise, we have to say that not each phage of this cluster revealed such a high level of identity to any other group member.

However, besides DSM 11031, all prophages had a counterpart within the cluster to bridge to the remaining representatives. The prophage DSM 11031 had only 94 % identity to some of the prophages, making it a new species per definition. Still, we keep this prophage within this group as we see the possibility of new isolates appearing, which can connect this phage to the remaining in the cluster. In addition, the majority of the investigated phage genomes originate from experimentally unverified prophages holding the risk of a 1 % gap, dividing prophage DSM_11031 from the rest. Due to the type strain SPβ within this cluster, it has already the ICTV approved species name *SPbeta*.

The majority of the *SPbeta* phages originate from host strains belonging to the *B. subtilis* species. The only deviating host is *B. vallismortis* DSM 11031 [CP026362], in which the prophage integrates into the *spoVK* locus and thus employs the same *attP/B* sites as other viral cluster representatives. The only *B. subtilis* prophage not in this cluster were from *B. subtilis* ATCC 13952 [NZ_CP009748] and *B. subtilis* J-5 [NZ_CP018295], both using *pbuX* as integration locus. To find out if those deviations base on false annotated host strains, we calculated average nucleotide identity values for all host strains (Figure 5 and Table S3). Results revealed *B. vallismortis* DSM 11031 with 91 % identity to be genomically close related to *B. subtilis*. Furthermore, *B. subtilis* ATCC 13952 proved to be a *B. amyloliquefciens* and *B. subtilis* J-5 a *B. velezensis* strains due to a high degree of average nucleotide identity to the remaining strains of the respective species clusters (Figure 5 and Table S3). Thus, the results show that phages of the *SPbeta* species prefer *B. subtilis* as host-strains. The second species cluster of the *Spbetavirus* contains 26 prophages. They are associated with phage H2 originating from *B. amyloliquefaciens* H (Zahler, Korman, Thomas, and Odebralski, 1987). For this reason, we propose this viral species be named *eta* for the Greek letter "H". All phages of that cluster associate with the host species *B. velezensis* and *B. amyloliquefaciens,* including those wrong classified *B. subtilis* strains mentioned above. Based on the genome data, *B. amyloliquefaciens* Y2 also has to be re-classified as *B. velezensis* (Figure 5).

**Figure 5:**
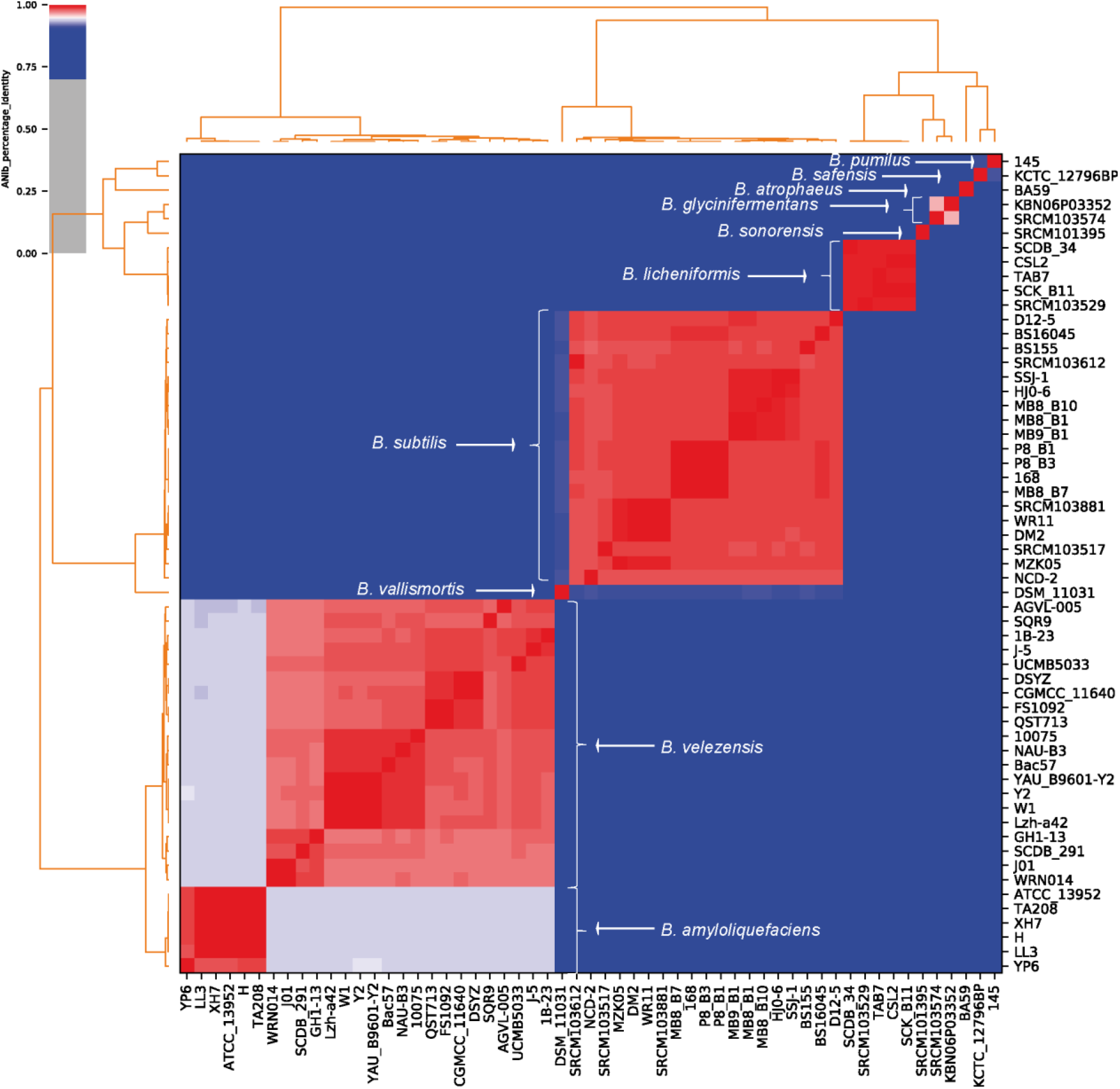
Average nucleotide identity analysis of SPβ-like host strains. Analysis reveals ten host species employed by Spbetavirus.

The next smaller viral cluster is formed by phages associated with *B. licheniformis.* The observed viral examples are the biggest among the identified *Spbetavirus*. Therefore, we propose this viral species be named *magnus*, which is Latin for big.

The following viral cluster is formed by phages associated with *B. glycinifermentans* and *B. sonorensis*. With 86 % identity (Table S3), both host species are related but still distinct. Therefore, this is the first viral species to be clearly associated with two host species. We propose these viral species be named *bimanducare*, which assembles from the Latin words *bi* meaning two and *manducare* for eating.

The remaining three phages 145, KCTC_12796BP, and BA59 are associated with *B. pumilus* 145, *B. safensis* KCTC_12796BP, and *B. atrophaeus* BA59, respectively. Like their hosts, these orphans present separate species. We propose no species names for those viruses yet and shift the naming until more representatives are discovered and investigated. For the first time, SPβ-like phages are classified taxonomically, revealing four species with several representatives within the genus *Spbetavirus* and three orphans with the potential to grow into independent new species. It is interesting to note, that the SPβ-like species narrowly associate with their host species implying a host-parasite co-evolution.

### What defines an SPβ-like phage?

Having 64 genomes of SPβ-like prophages "in our hands" gave us an excellent opportunity to explore the conserved proteins and functions defining this phage group. We performed a new open reading frame (ORF) calling with all prophage genomes to create an even starting point. The newly annotated proteins we used for orthology detection. Of the 928 identified distinct SPβ-like proteins, 25 were present in all prophages, and 13 were absent in one or two viruses (Table 2 and Table S4).

**Table 2:**
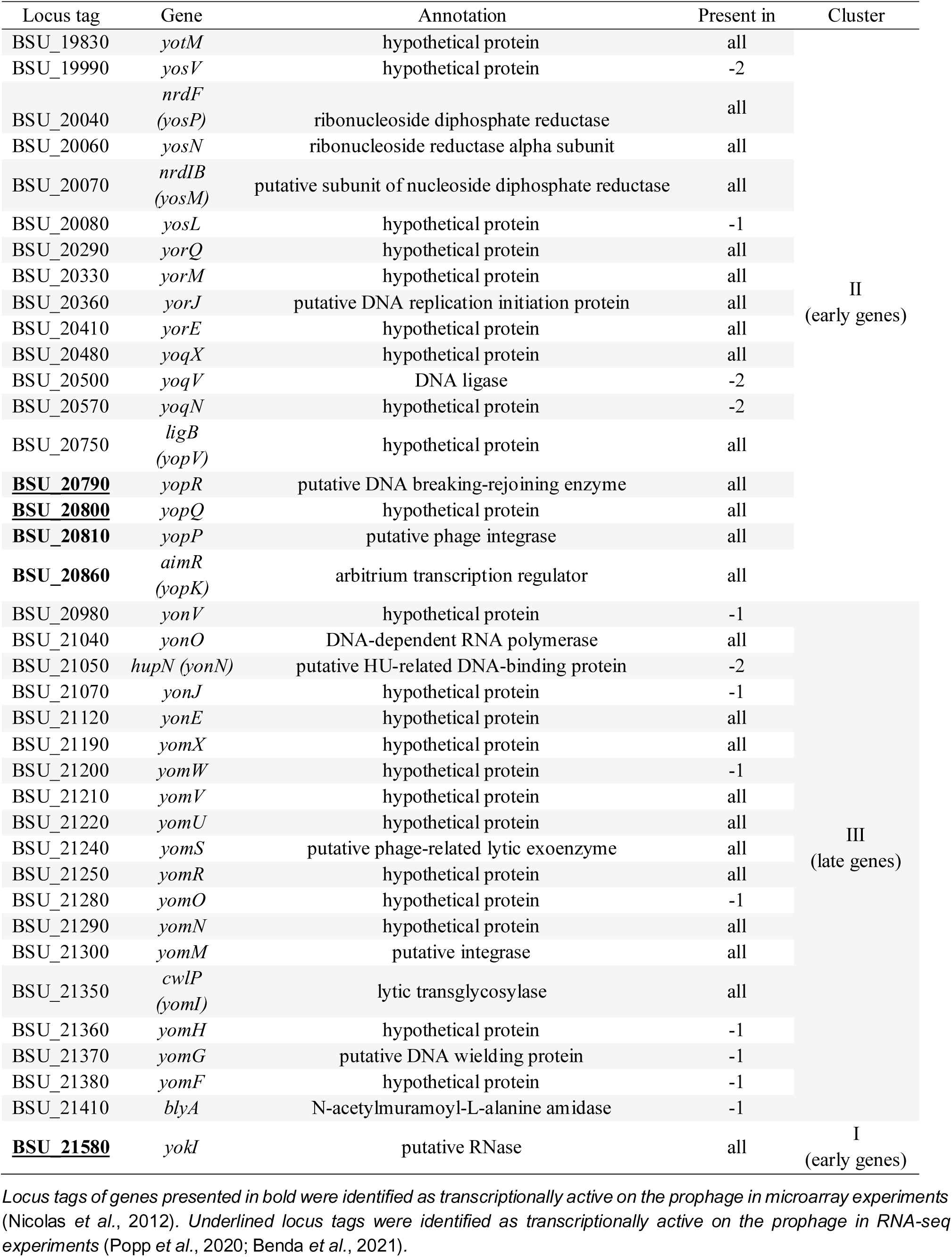
Core protein genes of the genus *Spbetavirus*

Individual prophages may be incomplete, like we assume it for *B. velezensis* 10075. It reveals 96 % identity to the prophage of *B. velezensis* Bac57 but has an about 17 kb smaller genome than Bac57 (Table 1). A big part of the missing genetic information consists of a 15 kb fragment in cluster III, containing conserved homologues from BSU_21360 to BSU_21480. It includes the *blyA* gene coding the phage lysin, a crucial component for a functional prophage (supplementary materials 6). Thus, we considered orthologues as conserved, even missing in one or two prophages, as long as they are present in our functionally verified wild type isolates.

The early cluster I, presenting about 20 % of the phage genome, contains just *yokI* as a conserved protein. In the latest annotation of *B. subtilis* 168 [NC_000964.3], *yokI* codes for a putative RNase [BSU_21580]. We assume this annotation is based on the PANTHER (Protein ANalysis THrough Evolutionary Relationships) Classification System (Mi *et al*., 2013). It was the only database we found to predict this function for YokI. However, this gene was also proposed to be the toxin of a type II toxin/antitoxin system (Van Melderen, 2010; Holberger *et al*., 2012). Our analysis may resolve this controversy. YokI is conserved in all SPβ-like phages but not its predicted counterpart, the antitoxin YokJ, only present in 20 of the 64 prophages. Obviously, YokI is not associated with its predicted antitoxin, making it unlikely to be a toxin itself. The existence of two *yokJ* deletion mutants, BKK21570 and BKE21570, further support this assumption (Koo *et al*., 2017). We propose YokI to fulfil an important function in the viral life cycle since it is conserved in all SPβ-like phages. However, this function is not essential under laboratory conditions. We could successfully delete this gene for the SPβ prophage without affecting its ability to generate viable particles (Figure 6 B III).

**Figure 6:**
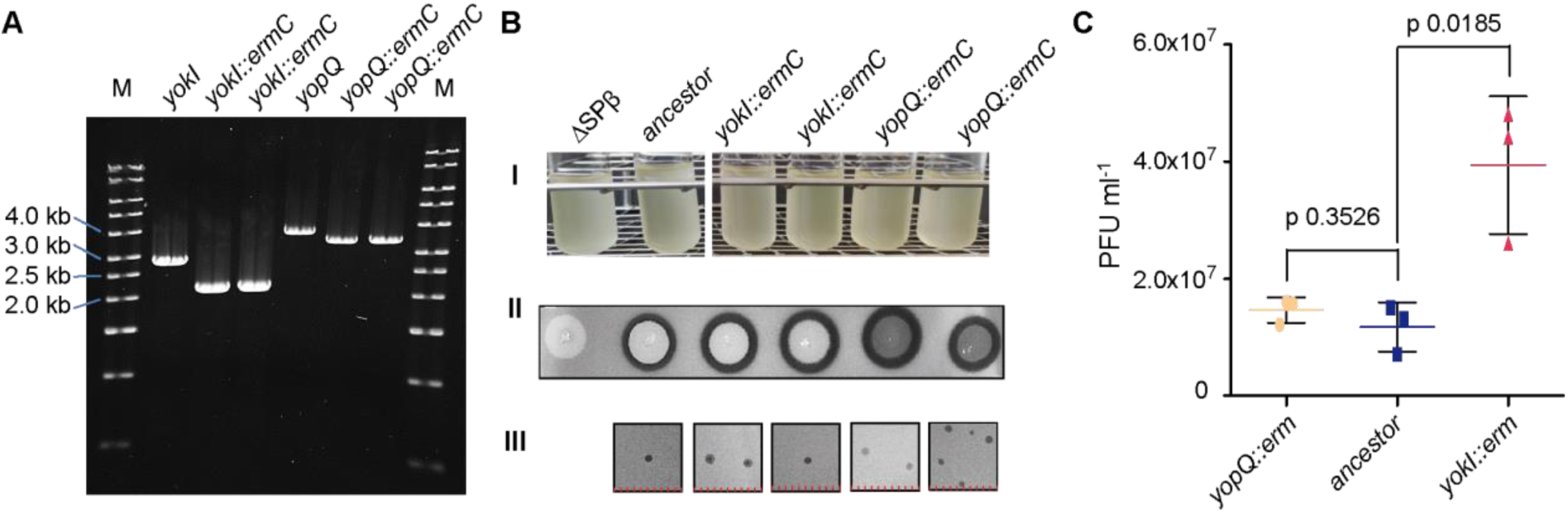
Investigation of the yokI, yopQ mutants. **A:** PCR control of the yokI and yopQ mutants. The primers PP342/PP345 were used to amplify the yokI locus and results with the wild type situation in a 2.7 kbp fragment and with the mutant in 2.1 kbp. To verify the yopQ locus, the primers PP073/PP319 were used. The wild type situation led to a 3.7 kbp fragment and the mutant to 3.5 kbp. All PCR fragments revealed the expected size. **B I:** Growth experiment of all investigated strains in liquid LB medium. No particularities were observed. **II.** Verification of SPβ prophages. Bacteria with an SPβ prophage secret sublancin 168, which is toxic to SPβ free strains. Lysogens applied to a layer of an SPβ free sensory strain generate an inhibition zone around themselves and thus verify the presence of an SPβ prophage in the tested strain. All tested mutant strains secreted sublancin 168 and thus proved the presence of SPβ. **III.** Verification of phage viability. All investigated mutants released viable virions in the supernatant confirmed via plaque formation. **C:** Release of viable SPβ virions into the supernatant in cultures of lysogenic bacteria. The SPβ yokI::ermC mutant releases about four times more virions compared to the ancestor strain of the mutant. The p-values were determined by an unpaired t-test. Horizontal bars indicate the respective mean values. Error bars indicate standard deviation.

The most conserved proteins, 19 to be precise, are located in the late cluster III, which should contain the structural genes of the phage (Table 2). This cluster also contains the *yonO* gene [BSU_21040], coding for a unique DNA-dependent RNA polymerase (RNAP) observed only with SPβ (Forrest *et al*., 2017), and the already mentioned *blyA* [BSU_21410] gene coding for an SPβ specific lysin (Regamey and Karamata, 1998). Those genes indicate the specific RNAP as a key feature of the SPβ phage-type, including specific structural components and a conserved SPβ-like lysin.

Early cluster II contains 18 conserved proteins (Table 2). Most of them are without functional annotation. Those which have an annotation indicate their involvement in the phage genome replication. This cluster’s most prominent and best-investigated gene is *aimR* coding for the arbitrium transcription regulator (Erez *et al*., 2017). The AimR protein is a key component of the decision-making system of SPβ-related phages, also known as the arbitrium system (Erez *et al*., 2017). It is part of the lysogeny-management system responsible for switching from lytic to lysogenic replication. Its presence among the core genes confirms the lysogeny-management system to be conserved among SPβ-related phages and implies some of the remaining homologs to be involved in lysogeny maintenance and resolvement. However, the earlier proposed lysogeny repressor YonR (Lazarevic *et al*., 1999) was only conserved in 15 prophages and thus not even in all phages associated with *B. subtilis.* Most important YonR was in none of the new wild type isolates, which are confirmed to form prophages.

We also did not identify the site-specific recombinase SprA to be conserved among the SPβ-like phages, even the presence of such a protein was set as an essential property during manually extraction and curations of the SPβ-like prophages. This contradicting observation shows that although SPβ-like phage all employ a site-specific recombinase, those are not all from the same type. In addition, the integrase type used by the phage correlates with its insertion locus. The 44 SprA orthologues proteins were just present in phages associated with *spsM*, *kamA*, *pbuX* and *spoVFB.* The remaining 20 used two other integrases and integrated into *spoVK* and *glnA*. The phage Goe14 contains such an integrase which prefers *spoVK* for integration. As mentioned before, that protein truncates itself upon prophage integration (Figure 1 C). It shortens its proteins sequence by 131 amino acids and creates a 97 amino acids long new ORF being completely homolog to the truncated C-terminal end of the phage-encoded integrase (supplementary materials 7).

A domain search of the Goe14 integrase revealed it to be of the same organisation as the SprA integrase of SPβ but with a prolonged C-terminus with no similarity to any known structure (supplementary materials 8). We compared the protein sequence of the Goe14 integrase to all proteins of *B. subtilis* 168, including SPβ. Surprisingly we did not observe any relevant similarity to SprA of SPβ but a pronounced one to SpoIVCA with 28.0 % identity (61.3 % similar). SpoIVCA is the site-specific recombinase from the *skin* element, a degenerated prophage of *B. subtilis* 168 (Abe *et al*., 2014).

The identified functional domains from Lzh-a42 integrase, which integrates the prophage into the *glnA* locus, were of distinct type and organisation (supplementary materials 8). Using its protein sequence as a query for a BLASTp search in the genome of *B. subtilis* 168 we could again reveal a related protein. It was the YdcL site-specific recombinase of the ICEBs1 element (Lee *et al*., 2007). Their relation is also underlined by the relatively large direct repeats produced as *attL* and *attR* upon integration of the alien genetic element. With the here investigated SPβ-like phages, it was 106 bp of the *glnA* gene and in the case of the ICEBs1 60 bp of the *trnS-leu2* gene [BSU_tRNA_51] (Lee *et al*., 2007). Additionally, the Lzh-a42 integrase is a tyrosine-type site-specific recombinase, while the integrases of SPβ and Goe14 are of the serine recombinase family.

These results prove the recombination unit of SPβ is a conserve core element of SPβ-like phages. Suzuki and colleagues (Suzuki *et al*., 2020) recently demonstrated that those are artificially interchangeable between *skin*, ICEBs1 and SPβ and still result in a functional SPβ derivate (Suzuki *et al*., 2020). Our *in silico* analysis indicate the interchangeability to be a frequent natural phenomenon. The best examples are Goe11 and Goe14. Both use different recombination units (Table 1) even they are gnomically very similar (Figure 4).

### Lysogeny management components of SPβ

To determine which conserved proteins could be the prophage management components, we evaluated the data of Koo and colleagues (Koo *et al*., 2017). To find the essential gene set, they individually addressed each gene of *B. subtilis* 168 with barcoded kanamycin (BKK) and erythromycin (BKE) deletion cassettes. This investigation also included the prophages of *B. subtilis* and thus also SPβ. Surprisingly, none of the SPβ genes was mentioned as essential, and we could recover BKK and BKE mutant numbers for all conserved SPβ genes. However, we noticed that not for all mutants barcodes were available. For instance, the *yopR* mutants have just a barcode for the erythromycin-based mutant [BKE20790] and non for the kanamycin-based mutant [BKK20790]. An opposite situation was observed for *yomS* [BSU_21240]. The *cwlP* gene [BSU_21350] is the only one not to have any barcode sequences at all, even two mutants were announced.

To determine which genes are transcribed from the dormant prophage, we analysed three independent transcriptome datasets from *B. subtilis* 168 under non-inducible conditions (Table S5). The first one from Nocolas and colleagues (Nicolas *et al*., 2012) was generated with a microarray technique. From this study, we only used the control data set of the mitomycin C induction experiments. Of the SPβ conserved genes, five genes were transcriptionally active, *yopR*, *yopQ*, *yopP*, *aimR* and *yokI* (Table S5). The second data set from Popp and colleagues (Popp *et al*., 2020) and the third data set from Benda and colleagues (Benda *et al*., 2021) were up-to-date Illumina RNA-seq data sets. They were generated as control experiments from cells cultured in LB-medium at 37 °C at vigorous shaking. Their results revealed *yopR*, *yopQ* and *yokI* to be transcriptionally active (Table S5).

To find out which of the three proteins may fulfil the function of a prophage repressor, we reproduced the deletion experiments of Koo end colleagues (Koo *et al*., 2017) on those specific genes. We successfully obtained *yopQ* and *yokI* mutants on an SPβ prophage (Figure 6 A). The *yopQ* mutants grew well in the liquid medium. However, they revealed a translucent structure on plates during the sublancin activity assays, employed to verify the presence of the prophage (Figure 6 B). This observation implies cell lysis during the stationary phase. The *yokI* mutants revealed no noticeable difference in colony morphology but released about four times more SPβ virions into the supernatant than its ancestor strain (Figure 6 C).

We struggled to obtain mutants of the *yopR* gene, implying it to be the SPβ repressor. Potential *yopR* prophage mutants appeared about two days after transformation in low numbers, and their genotype could not be confirmed. Just recently and independent from our investigation, Brady and colleagues (Brady *et al*., 2021) experienced the same struggles during their *yopR* deletion attempts. Indeed, their results revealed YopR as the master repressor of the SPβ prophage (Brady *et al*., 2021). Our concept to prove the hypothesis of YopR being the SPβ repressor was to isolate clear-plaque mutants and investigate their genome. We isolated six of these mutants cpm1 to cpm6 from the supernatant of *B. subtilis* 168 and verified their genomes for consistency via PCR. Two of six might have faced deletions in their early operons (supplementary materials 9). Such deletions were reported in the literature for earlier identified clear-plaque mutants of SPβ (Spancake and Hemphill, 1985). The cmp2 mutant with a potentially complete genome was whole-genome sequenced with Illumina technology. Using the obtained reads for single nucleotide polymorphisms (SNPs) analysis, a deletion of 16 bp became evident in the *yopR* gene and a subsequent frameshift truncating the coding region. Next, we PCR amplified, and Sanger sequenced the *yopR* genes of the remaining clear-plaque mutants cmp3, 5 and 6 with a potentially complete genome. We observed one deleted base and one point-mutation in cpm6, which likewise in cpm2 lead to a frameshift in *yopR* and truncation of the gene. In both cpm3 and cpm5, we observed deletions of the entire intergenic region of 23 bp between *yopQ* and *yopR* (Figure 7).

**Figure 7:**
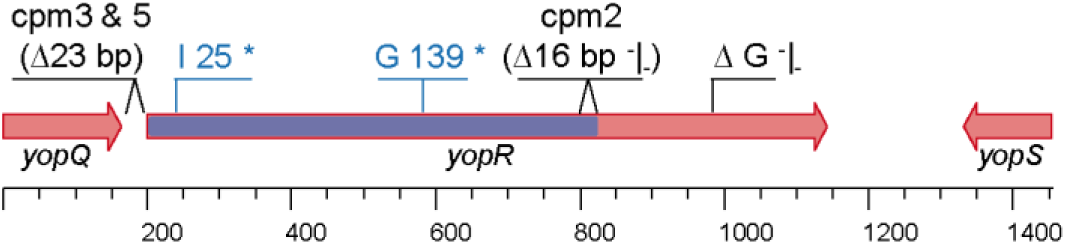
Mutation analysis of yopR clear plaque mutants (cpm). Mutations indicated in black were discovered during the presented study. Mutations presented in blue were discovered by Brady et. al. (Brady et al., 2021). The bar in the yopR gene marks the predicted DNA breaking-rejoining catalytic core domain. * = stop codon. ^-^|- = frameshift.

All observed mutations affected the 325 amino acids (aa) long YopR protein. The observed deletion of the entire *yopQ* and *yopR* intergenic region removed the ribosome binding site of *yopR* and thereby likely abolished its translation. To find out the impact of the remaining mutations, we scanned YopR for the presence of known functional domains. Using InterProScan5 we could identify a predicted DNA breaking-re-joining catalytic core between 12 to 218 aa. Interestingly, only the 16 bp deletion of cpm2 affected the last six C-terminal amino acids of the predicted domain. The point-mutation of cpm6 was far behind at amino acid 273. To test whether these *yopR* mutations are responsible for the observed clear-plaque phenotypes, we created a host strain with an artificially expressed YopR. Therefore, we fused *yopR* with the constitutively active P*_alf4_* promoter and introduced it into the *amyE* locus of the *B. subtilis* TS01 chromosome. Employing the so created strain as host, clear-plaque mutants cpm2 and cpm6 lost their lytic phenotype. They could not form plaques anymore (supplementary materials 10). Consequently, our results are in good agreement with the observations by Brady and colleagues (Brady *et al*., 2021) and confirm YopR to be the prophage repressor of SPβ.

## Discussion

### Genome replication and packaging

Our investigation revealed SPβ-like phages to replicate their genomes in a concatemeric way and pack their genomic DNA via the head-full mechanism into their proheads. Such a packaging system is known from the *Escherichia virus* P1. The concatemeric viral genome is pumped into its prohead until it is full. This triggers an unspecific cut of the DNA and the release of the translocase machinery, which still holds the genome. An association with a new prohead initiates its filling with the remaining viral genomic material (Calendar, 2006).

SPβ presents itself as not very efficient in this type of genome packaging. Its apparent inability to continuously package its genome concatemer may be the reason for the relatively low replication rate (Figure 2 A). Restarting genome translocation at the pac-site after each 100 %-plus filled phage head would be an enormous waste, as it would leave a < 100 % fragment with each 100 %-plus filled phage head. Even if the left behind fragments were sufficient to ensure phage reproduction, one way or another, they still would be doomed to degeneration due to the missing pac-site, which was packed as the "plus" with the preceding filling. A phage like Goe11, which is much more successful in its reproduction judging by its plaque size (Figure 1 B), could draw its advantage from the more efficient use of its genome concatemer. If we hypothetically assume the entire reproductive process of SPβ and Goe11 to be the same, except for the genome packaging, Goe11 would generate about twice as much viable offspring as SPβ just by the continuous use of the genome concatemer. This efficient use of Goe11’s genome concatemer is evident in the sequence coverage plot from its particle-packaged genomic DNA (Figure 2 C). To prove this hypothesis, one could replace the genome packaging components of SPβ with those of Goe11 and investigate the plaques and offspring numbers of the so produced hybrids. Before we can do so, we first need to identify and experimentally verify the genome packaging machinery of SPβ.

All in all, these data lead to the conclusion that SPβ-like phages pack their genomes with a head-full mechanism placing 100 %-plus genome per virion. The question of what the "plus" could be, cannot be precisely answered from the available data. The frequently observe ∼250 bp block after the opening point unlikely represents the genome overhang (Figures 2, right panel). It is precise and equal in size with all investigated isolates, even though the isolates reveal distinct genome sizes and thus hold the potential to pump overhang fragments of different sizes into their viral heads. We can speculate that the 250 bp fragment is associated with a very likely conserved translocation machinery and contains the pac-site sequence pattern. All packaging start points were the same for the investigated phages (supplementary materials 11).

The Goe14 mapping reveals a focused read accumulation of about 2 kbp following the ∼250 bp peak (Figure 2 D). It represents 1.6 % of the whole genome and could be thought of as the "plus" overhang. The *Bacillus* phage SPP1, a smaller lytic *Syphoviridae* type of phage, also employ pac-sites and a head-full genome packaging mechanism (Calendar, 2006). With 1.4 kb it reveals similar terminal redundancy with the particle packed genomic DNA (Camacho *et al*., 2003). However, for SPβ-like phages, this assumption has to be experimentally verified in the future. The observed sequence accumulation of ∼2 kbp was only observed with Goe14, and its proportions are not balanced to the average coverage. It should only be twice as high as the rest of the coverage if the observed accumulation originates from the above-average initiation of genome translocation at the pac-site and the 100 %-plus head-full mechanism. However, here it is significantly higher (Figure 2 D).

### Expansion of the genus *Spbetavirus* and their prophages

Isolation of new SPβ-like viruses, particularly Goe14, brought us on the track of new prophages. Their isolation revealed a whole new diversity of SPβ-related phages. For the first time, new species of the genus *Spbetavirus* were identified, strongly associated with their host genus.

We could identify five more SPβ prophage integration loci, of which *spoVK* is also actively used in derivates of *B. subtilis* 168, and *glnA* at least was used in the past by a potentially related SPβ derivate. It remains unclear if the other three integration loci, present in *B. subtilis*, are actively targeted for prophage integration. The only two potential *B. subtilis* lysogens using *pbuX* for prophage integration revealed themselves to be of a distinct bacterial species (Table 1). However, the phage H2 is known to replicate and lysogenize *B. subtilis* CU1050, an SPβ free derivate of 168 (Zahler, Korman, Thomas, Fink, *et al*., 1987). That observation suggests the possibility of *pbuX* as an integration locus in *B. subtilis*. Ultimately, this question can only be clarified by experimental studies.

From φ3T and SPβ, it is known that *spsM* and *kamA* are re-established during sporulation. How it appears with the here identified new integration loci is to be investigated. However, the *spoVFB* gene from *B. weihenstephanensis* KBAB4, which is disrupted by a prophage-like element, is re-established in the mother cell during spore formation to ensure dipicolinic acid synthetase (Abe *et al*., 2013).

During sporulation of *Geobacillus thermoglucosidasius* C56-YS93 a similar gene reestablishment and excision of a prophage-like element was observed with the *spoVR* gene (Abe *et al*., 2013). Its gene product is required for spore cortex formation (Beall and Moran, 1994).

It cannot be answered at this point if there are more potential integration loci for SPβ-like phages. However, this is likely to be the case since other sporulation-related genes like *cotJC*, *gerE*, *yaaH* and *yhbH* are frequently used for phage or transposon related integrations (Abe *et al*., 2013). Although, the mentioned examples were observed in bacterial species, not of the Subtilis-clade. Still, *spoVFB* was first observed as integration locus of the prophage-like element *vfbin* in *B. weihenstephanensis* KBAB4 (Abe *et al*., 2013) and now revealed itself as an integration loci of a *Spbetavirus* species (Table 1). The fact that SPβ-like phage can interchange their recombination units makes us expect further diversity.

BLASTn based prophage predictions were used as a starting point for the investigation of SPβ prophage diversity. The genome of the type-strain SPβ served as a query. Thus, it is likely that just very related prophages were identified in closely related host strains (Figure 4 and 5). Most successfully extracted SPβ-like prophages originated from *B. subtilis* genomes and genomes of related *B. velezensis* and *B. amyloliquefaciens* strains. Only a few prophages came from more distinct bacterial strains. Phages on the genus border with slightly over 70 % average nucleotide identity to the remaining isolates came from *B. safensis* and *B. pumilus* (Figure 4). If repeating the prophage search with those unique phage genomes as a new query, new and more diverse *Spbetavirus* members could be identified. Each round of further repetitions could even reveal the presence of SPβ-like phages ahead the Subtilis-clade. However, as further search attempts would base on experimentally not as functionally verified prophages, the resulting error multiplication can hardly be estimated. Thus, it is better to start after prophage 145 from *B. pumilus* and KCTC_12796BP from *B. safensis* are verified functional.

### Multiple lysogenisation

Having so many phages associated with one host species like *B. subtilis,* leads to the question of how many phages can simultaneously lysogenise one host. For SPβ and φ3T it is known they can lysogenize the same host (Warner *et al*., 1977). To our surprise, we frequently observed multiple fragmented SPβ-like phages in one strain. For example, *B. velezensis* DKU_NT_04 [NZ_CP026533] has rudiments of prophages in three of six insertion sites. An approximately 19 kbp fragment sits in *spsM*, an approximately 262 kbp fragment sits in *glnA*, and a not further defined fragment in *pbuX*. This strain suggests that multiple lysogenization beyond two phages may be possible. However, we never observed more than one complete SPβ-like phage in the same host strain. One can only speculate about the reasons. It is conceivable that phages from the same genus are simply too similar to persist in the same host. Their similar genomic organisation and high sequence homology make recombination of the prophages likely. That can lead to the degeneration of both involved entities or to the formation of hybrids.

A possible indication for the degenerative recombination hypothesis is provided by the SPβ-like prophages of *B. velezensis* SGAir0473. Here, we identified a prophage in both the *spoVK* and *spsM* genes. Both seem to be fragmented. The prophage in *spoVK* is too large for an SPβ-like phage with the size of ∼ 205 kbp, and the prophage in the *spsM* gene is too tiny with ∼ 30 kbp. However, the total size of ∼ 235 kb is about the size of two intact representatives. This example implies that the simultaneous presence of two SPβ-like phages represents an unstable structure. Dragoš and colleagues were able to observe the formation of possible hybrid phages and even verified them as stable viral entities (Dragoš *et al*., 2021). However, we also observed assembly artefacts during an automated genome assembly of an SPβ and φ3T double lysogenic strain (unpublished data). Suppose the focus of such a genome assembly is not on the phages. In that case, this circumstance can easily be overlooked and lead to artificial genomes.

Finally, although the genomic study of diverse lysogens did not reveal any unambiguous double lysogens, we can suggest that this phenomenon is not limited to SPβ and φ3T. It may even have a biological function, such as creating new phage types (Dragoš *et al*., 2021). It remains an exciting question which factors allow such a double lysogen and which role the phage repressor plays since a superinfection of the same phage in a lysogen is not possible.

### SPβ core genes and its lysogeny management system

The identified conserved SPβ proteins are mainly without functional annotation. But even so, they can help to check new isolates for their relationship to SPβ if the nucleotide sequence differs significantly. Aligning the core genome to existing SPβ prophage transcriptomes proved to be a valid strategy for the identification of the SPβ lysogeny management components. This strategy is particularly interesting for other bigger prophages like SPβ, which bring several accessory genes relevant for the host and actively transcribed from the dormant prophage (Hemphill *et al*., 1980; Dragoš *et al*., 2020) (Table S5).

YopR is not the first protein proposed to be the prophage repressor of SPβ (Brady *et al*., 2021). Earlier, the YonR was proposed to be the lysogenic repressor of SPβ due to its similarity to the lysogenic repressor Xre for lysogeny maintenance of PBSX and YqaE of the *skin* element (Lazarevic *et al*., 1999). The d-protein (YomJ) also conveyed resistance against SPβ and closely related phage strains if expressed ectopically from a plasmid (McLaughlin *et al*., 1986). However, neither *yonR* nor *yomJ* are conserved (Table 2), and both could be individually deleted from the prophage genome without activating the prophage (Koo *et al*., 2017). However, genomic investigation of our clear plaque mutants confirmed YopR as essential for prophage establishment. Combining our results, we can even extract more knowledge about YopR. This protein has a "DNA breaking-re-joining enzymatic core" which Brady and colleagues propose as essential for the function of YopR as a repressor (Brady *et al*., 2021). In fact, the two clear-plaque mutants described by the group had their mutation in this particular region. The mutations of the clear-plaque mutants we identified lie at the end or behind the predicted domain (Figure 7). Particularly cpm6 proves YopR to contain new, previously unknown domains to explore.

Besides *yopR* there were two more conserved genes transcribed from the prophage. We could successfully delete both genes without induction of the prophage. The translucent cells of the *yopQ* mutants and the significantly increased spontaneous release of viable virions of the *yokI* mutants imply both proteins to be involved in the lysogeny management system of SPβ (Figure 6). The phenotype of the *yopQ* mutants does not directly allow this conclusion, but since this gene forms an operon with *yopR* and is thus transcribed together with it, we assume its involvement in lysogeny management. The phenotype of the *yokI* mutant is obvious. In Figure 8 we present our hypothesis about how YokI might be connected with the lysogeny management system of SPβ.

**Figure 8:**
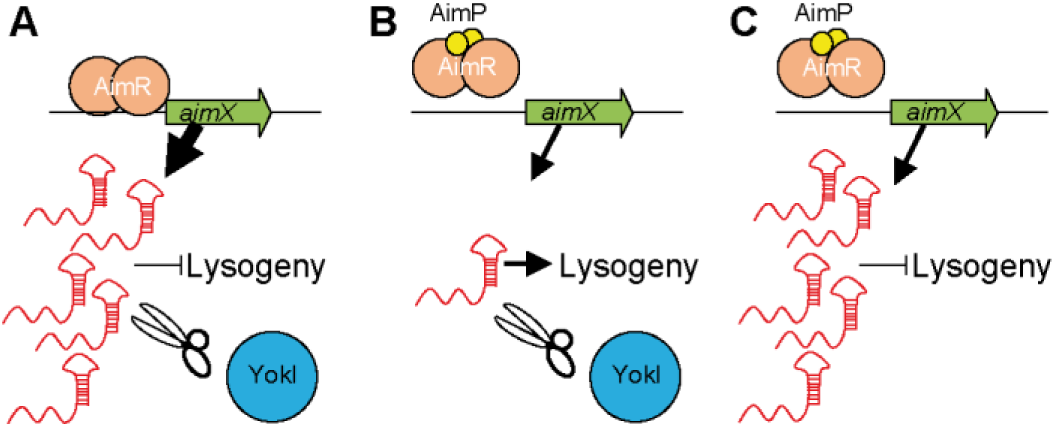
The hypothetical role of YokI as AimX concentration balancer. **A:** Early infection situation. AimR binds to the aimX promoter and activates transcription of the ncRNA AimX, which in turn represses lysogeny. YokI degrades the constantly transcribed AimX and avoids AimX accumulation. **B:** Late infection situation. The signal peptide AimP binds to AimR, the complex dissociates from the aimX promoter and AimX transcription is reduced, YokI degrades the remaining AimX what activates lysogeny. **C:** Late infection situation in deletion yokI mutant. The signal peptide AimP binds to AimR, the complex dissociates from the aimX promoter reducing AimX transcription. However, in the absence of YokI, present AimX is not degraded, accumulates, and reduces lysogeny.

The arbitrium system consists of AimR, a transcription activator of the non-coding small RNA (ncRNA) AimX, and a quorum-sensing signal peptide AimP. When AimP is processed and re-imported into the cytosol, it interacts with the AimR dimer. Subsequently, this complex experiences a conformational change, dissociates from the aimX promoter, and reduces *aimX* transcription (Erez *et al*., 2017). Brady and colleagues showed that *aimR* mutants mainly prefer lysogeny and *aimP* mutants the lytic route (Brady *et al*., 2021). Thus, *aimX* transcription is reduced without AimR and increased without AimP. The AimX ncRNA may modulate the function of YopR and make it more slippery or tight. However, the arbitrium system is not essential for lysogeny establishment or resolvement but rather fine-tunes the system (Erez *et al*., 2017; Otte *et al*., 2020; Aframian *et al*., 2021; Brady *et al*., 2021). The question not asked before is what happens with the ncRNA AimX itself. As a strongly transcribed ncRNA it needs to have an equilibrium mechanism. Something which ensures this ncRNA does not accumulate, like an RNase keeping it in balance. YokI is the perfect candidate. It is conserved in the *Spbetavirus*, transcribed from the inactive prophage, and contains a predicted RNase domain. The increased amount of virus particles released into the supernatant of *yokI* mutants could be explained by the accumulation of AimX. In such a case, it would no longer be counterbalanced and accumulate like in the AimP mutant and push the system to lysis (Figur 8). This hypothesis is also supported by historical data where lost fragments between *sprA* (*yokA*) and *sunI* (*yolF*), including *yokI*, were associated with unstable lysogeny (Spancake and Hemphill, 1985).

To explore the *yokI* hypothesis and many other aspects of SPβ revealed during this work, we will go ahead of the genomic perspective and devote ourselves to specific experiments to further unravel the mysteries of SPβ.

### Data Availability

Phage genome sequence data used for the presented investigation were either obtained from SRA and GenBank repositories or created during this investigation and subsequently submitted to the public SRA and GenBank repositories. The respective accession numbers can be found in Table 1 and in supplementary materials 1.

### Supplementary data

Supplementary data are available at bioRxiv online.

### Funding

This work was supported by Volkswagen Foundation [Re. 94045] and the Max Buchner Research Foundation [Re. 3799]. A.D. and V.A.F. were supported by Slovenian Research Agency [N1-0177].

## Acknowledgement

This work was also supported by the Georg-August University of Göttingen and the Brandenburg Technische Universität Cottbus-Senftenberg. We thank all FG Synthetic Microbiology group members for stimulating discussions and suggestions.

