## Supplementary_materials for "The *Bacillus* phage SPβ and its relatives: A temperate phage model system reveals new strains, species, prophage integration loci, conserved proteins and lysogeny management components"

### 1. Sequence data

#### Genome sequence data

| Strain | Data type | Accession number | Reference |
|---|---|---|---|
| SPβ cmp2 | Illumina 150bp paired | PRJNA775004 | this study |
| φ3Ts | Illumina 300bp paired | SRR11587866 | (1) |
| vB_BsuS-Goe11 | Illumina 300bp paired | PRJNA547503 | this study |
| vB_BsuS-Goe12 | Illumina 300bp paired | PRJNA547504 | this study |
| vB_BsuS-Goe13 | Illumina 300bp paired | PRJNA547505 | this study |
| vB_BsuS-Goe14 | Illumina 300bp paired | PRJNA774947 | this study |

#### Transcriptome sequence data

| Strain | Data type | Accession number | Reference |
|---|---|---|---|
| *B. subtilis* 168 | Illumina 150bp unpaired | SRR10487157 | (2) |
| *B. subtilis* 168 | Illumina 150bp unpaired | SRR10487158 | (2) |
| *B. subtilis* 168 | Illumina 150bp unpaired | SRR10487159 | (2) |
| *B. subtilis* 168 | Illumina 150bp unpaired | SRR12349871 | (3) |
| *B. subtilis* 168 | Illumina 150bp unpaired | SRR12349872 | (3) |
| *B. subtilis* 168 | Illumina 150bp unpaired | SRR12349873 | (3) |

#### 2. Goe14 host phage chimaera sequence read

>M03741:272:000000000-CPR7B:1:1117:18247:17205

GTACGGGAAAAACGGCGGTTGCACGGCTGAGCGGCAGGCTGTTCTTTGAAATGAATGTCCAGTCAAAAGGCG
GCTTAATAGAAGCGGGGCGGGCGGAGCTGGTCGAGGAGTATATCGGACACACCGCCGAAAAGACGAGAGAT
TTAATCAAAGAGTCATTAGGTGGCATTTTGTTTATAGACGAAGCCTACTCGCTGGCAAGAGGCGGCGAAAAGG
ACTTTGGGAAAGAAGCGATCGACACACTTGTAAAGAACAAGGTTTTAACTTTTATGCAACTTCTCAACTCGATA
TCCCTCAGCCT

*B. subtilis* yellow

Goe14 blue

*attL* green

#### 3. Primer used in this investigation

| Name | Sequence 5' | purpose |
|---|---|---|
| PP316 | CCTTCCTAGGCTAAAGAGTG | verification *yorN* |
| PP317 | TGATGTGGGCATCCATATCG | verification *yorN* |
| PP318 | AGCCTAGACGAGTTGGAAAG | verification *yopR* |
| PP319 | GCAGCTGAGCGACTATAATC | verification *yopR and yopQ* |
| PP320 | CGTGGTGGAAGTGGGAGATG | verification *yonJ* |
| PP321 | CAACGCCTATCCGACTTATC | verification *yonJ* |
| PP322 | TGTCGTGTTGCCTTTGACAG | verification *yomI* |
| PP323 | TCCCTCCAAGGAGATATCAG | verification *yomI* |
| PP324 | ATTGCCGGATCTGTTTGACC | verification *yolE* |
| PP325 | CAATTCCGTGAGCATGGTAG | verification *yolE* |
| PP312 | GGAACTAAGGTCGCAAATGG | verification *yokI* |
| PP326 | GTCCCATAAGCCGGTTACAG | verification *yokI* |
| PP327 | CAAGTACGGTCATCCCTTTC | verification *yokE* |
| PP328 | AAGGATTGGCCAGAGTAGCG | verification *yokE* |
| PP359 | TGTTTACCGGTGGATCGATG | verification *yosL* |
| PP360 | CTATGTACGGCCTCCTTATC | verification *yosL* |
| PP073 | tacgTGATGCCTTCATCAACTAGA | verification *yopQ* |
| PP342 | TTTCACCGGACAAAGCAACC | verfification *yok*I |
| PP345 | CCGCATTGCACATCTTTAGC | verfification *yok*I |
| PP375 | aaa*GAATTC*ttgtcaagtgaaggcgcgctatgctacaatacagcttggtttaa**aggagg**aaacaatcATGTTCAATAGTGAGATTAAGGAA | cloning *yopR* (EcoRI) |
| PP376 | ttt*GGATCC*attacatgat**cctcct**taTTAAATGGTCGTCTCTTTTAGAC | cloning *yopR* (BamHI) |

The sequence in capital letters corresponds to the region to be amplified. The sequence represented by lowercase letters represents an artificially attached overhang. Italic capital letters represent attached restriction sites. The sequence shown in bold lowercase letters corresponds to an attached ribosome binding site.

#### 4. Plasmids used in this investigation

| Name | Relevant genotype | Reference |
|---|---|---|
| pAC7 | *amyE'-aphA3-lacZ-'amyE* | (4) |
| pRH167 | *amyE'-aphA3-* $P_{afl4}$*-yopR-lacZ-'amyE* | this study |

The $P_{afl4}$-promoter is an artificially designed constitutively active promoter (5).

#### 5. Bacterial strains used in this investigation

| Strain | Relevant genotype | Reference |
|---|---|---|
| *E. coli* DH10B | str. K-12 $F^-$ Δ(*ara-leu*)*7697*[Δ(*rapA'-cra'* )] Δ(*lac*)*X74*[Δ('*yahH-mhpE*)] duplication(514341-627601)[*nmpC-gltI*] *galK16 galE15* e14$^-$(*icd*$^{WT}$ *mcrA*) ϕ80d*lacZΔM15 recA1 relA1 endA1* Tn*10.10 nupG rpsL150*($Str^R$) *rph*$^+$ *spoT1 Δ(mrr-hsdRMS-mcrBC)* $\lambda^-$ Missense(*dnaA glmS glyQ lpxK mreC murA*) Nonsense(*chiA gatZ fhuA*? *yigA ygcG*) Frameshift(*flhC mglA fruB*) | (6) |
| *B. subtilis* 168 | trpC2 | (7, 8) |
| *B. subtilis* Δ6 | trpC2; ΔSPβ; subclacin 168-sensitive; Δskin; ΔPBSX; Δprophage 1; Δpks::Cm; Δprophage 3; ΔICEBs1 | (9) |
| *B. subtilis* TS01 | Δ6; *amyE'-ermD-* $P_{mtlA}$*-comKS-'amyE* | (10) |
| *B. subtilis* BKE20790 | 168; *yopR*::*ermC* | (11) |
| *B. subtilis* BKE20800 | 168; *yopQ*::*ermC* | (11) |
| *B. subtilis* BKE21580 | 168; *yokI*::*ermC* | (11) |
| *B. subtilis* KK001 | TS01; SPβ c2 | this study |
| *B. subtilis* KK002 | TS01; Δ*ermD*; SPβ c2 | this study |
| *B. subtilis* KK003 | Δ6; *amyE'-aphA3-'lacZ-'amyE* | this study |
| *B. subtilis* KK004 | Δ6; *amyE'-aphA3-* $P_{afl4}$*-yopR-lacZ-'amyE* | this study |
| *B. subtilis* KK005 | TS01; Δ*ermD*; SPβ c2, *yokI::ermC* | this study |
| *B. subtilis* KK006 | TS01; Δ*ermD*; SPβ c2, *yokI::ermC* | this study |
| *B. subtilis* KK007 | TS01; Δ*ermD*; SPβ c2, *yopQ::ermC* | this study |
| *B. subtilis* KK008 | TS01; Δ*ermD*; SPβ c2, *yopQ::ermC* | this study |

#### 6. Genome comparison of phage 10075 to Bac57 and SPβ

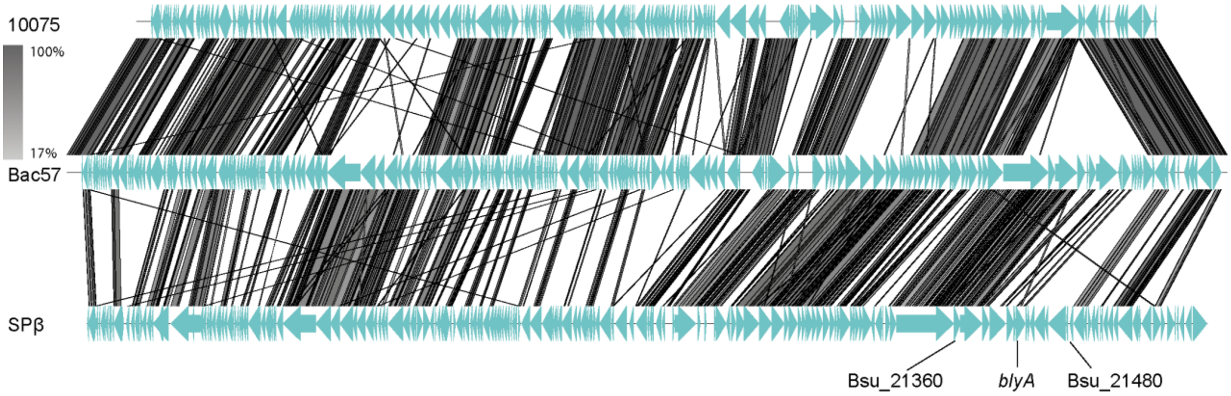

Genomes were compared using EasyFig 2.2.5 with the tblastx algorithm. Phage 10075 lacks the corresponding region of SPβ between gene Bsu_21360 and Bsu_21480, including the *blyA* gene coding the phage lysin.

#### 7. Goe 14 integrase

The region of the N-terminal part, present in the circular and prophage form, is highlighted green. The C-terminal protein part, which forms a new small hypothetical protein after prophage integration, is highlighted yellow.

##### Goe14 integrase in the phage genome

>Goe14 integrase

TTGCTTATATTAAAGGGAGATCAAAAGCTAAGTACGGACAATATAATGCATTTGATTGGCAAAATTAATGCATT
GATTTATGCCAGAGTATCAACAACAGATCAAGCTAATAAGGGTTTTTCAATTGAGTCACAAATAGAAAGGTGT
AAAGAAAGAGCAATCTCTAAATTTGGATACAAGGAAAGCGAAATAATTGCGTTAGTTGAACCGGGGGGCATG
GGAGATGATCCAAACCGGCCAGCTCTTAATCACGCCTTATATTTATTAGAAAAAGGACTAGGAAAAAAATTCAT
AGTATTACACCCAGATCGTCTAACTAGAGATAACACCTTACAGGGTGTGGTATCAAGAAGAATATGGGGTATG
GGTGTTGATATCGAGTTTATTGAATTTGAAGTAAATCCTCATGATCCTGAATCAATGCTGATGTACAACATTCA
AGGTTCAATTGCACAGTACAATAAGGCGAAAATTCATGCTAACTCAAAGCGAGGAAGATTGGCTAAAGCAAA
GAAAGGTGAATTTCCTTCATTTAAAAGGTTGTATGGGTACAAATTTAATACAGTGACAGATCTTCCTGAGTATA
ACGAAGAAGAAAAAGAAATTTTGCTTGAAATGAAGGATATGCTCTTAAACAAAAAAATGTCCTCGAATGAAAT
TGCTAAAGAGCTTTCAAGAAGGGGAGTTGCTGCTCCTAACGGAAAGACTTGGTATCAGGCAACTGTAAGTAG
AATGTTGCAGAATGAAGATTACACAGGTGATTTTTATTATGGAAAGTCAAAAGTTGTTCAAATTAATGGTAAGA
AGAAACAGGTGCCGACAAACAAAGACGAATGGATATTGATAAAGATTCCTCCAATGTGGGATAAAGCTACAC
GAGAGCAAATCATTGAGCGATTGAAGAGCAACTTTAAGGGAAGAAGTCGAGCTACTAAAGACTATTTGCTGA
AAAACAAGGCGAAATGTGGTCGTTGTGGAGGGGCATGTGGCTCAGGAATAACTTCTAAAACTAAATCAGGCG
TATATAAGTATTATTCTTGTAGAGCAAAGACTGCAAAAGGATATCAAAATGGCAAGAAAGTTGTTTCATGTGA
AGGTAAGAACTGGAGAGTCGATATCGTTGATGAAGTTTTTTGGAACTGGTTTATAAAACTCCTGAAGAATCCA
GAGAAATTTCTAGAATCATTTTTAGAGGAAGCGTCTGATCAAAAGAAAATCGATGAACTCAAAGCAAAAGCAA
GTCGACTAGAAAAACAACTCGGTGAGATAGATGAAGAAATAGCAAATTATGTAATTCTTTTTGGGAAAGGGAA
AATTAAAGAAAGTATGTTCGACCAACTGTCTCAGCCATTGGAACAGAACAAAGAGCATATTGAGAATGAGATT
AAAATAATTAATTCACAATTGGCTGCAAATAAACAAACTGAAGACAAAAAACAAAAAATGATCGAGTATATTG
GCTCGTTCTCTAAAATGATTAAAAATGAAATTACAATAGAAGAAAAACGGCAATTCCTTGATTTCTTCATTGAA
AAAGTAACTCTGTTTGATGATGATCACATGGAAGTTGTATGGAAAAGTTCTTCACTTAATAATGAAAACAGCCA
TGTGGATTTTCTTAACGGCGAGCAGGGAGGGGAGTTTATAAACTCGCATAAAAGACTAAACCATATTCAAGAA
CAAGGTTTTAACTTTTATGCCACTTCTCAACTCGATATCCCTCAGCCTCAACGTCAGCATAAATACGTATGGCAG
CAATATTTAGAGCAATTTAATGAGATCAAAAAAATGCACTTTGAAGAGCTTAAAACTGTAGGAGAGATTTCTAA
GGAACTAAATATCTCAGATTGGATTATTTTAGATTTATTTAAATCACAAAATGTGGACAAGCTTTCATTTCAGGA
ATTGTCTAAGCGTAGACGAACTAAGGACTTTGCTTTTCTTTATGATCTGCATTTCAATAAGAAAATGAGCCTGA
AGGAAATTAGTCGAGCTTTCGATTATTCTCCACCTTATATCCGTCAAGTTTTTAAGGATCAGGGAATTAAACACT
TAACCTTTAAAAATCAGTATAAAAATTAA

>Goe14 integrase

LLILKGDQKLSTDNIMHLIGKINALIYARVSTTDQANKGFSIESQIERCKERAISKFGYKESEIIALVEPGGMGDDPNRP
ALNHALYLLEKGLGKKFIVLHPDRLTRDNTLQGVVSRRIWGMGVDIEFIEFEVNPHDPESMLMYNIQGSIAQYNKAK
IHANSKRGRLAKAKKGEFPSFKRLYGYKFNTVTDLPEYNEEEKEILLEMKDMLLNKKMSSNEIAKELSRRGVAAPNGK
TWYQATVSRMLQNEDYTGDFYYGKSKVVQINGKKKQVPTNKDEWILIKIPPMWDKATREQIIERLKSNFKGRSRAT
KDYLLKNKAKCGRCGGACGSGITSKTKSGVYKYYSCRAKTAKGYQNGKKVVSCEGKNWRVDIVDEVFWNWFIKLL
KNPEKFLESFLEEASDQKKIDELKAKASRLEKQLGEIDEEIANYVILFGKGKIKESMFDQLSQPLEQNKEHIENEIKIINS
QLAANKQTEDKKQKMIEYIGSFSKMIKNEITIEEKRQFLDFFIEKVTLFDDDHMEVVWKSSSLNNENSHVDFLNGEQ
GGEFINSHKRLNHIQEQGFNFYATSQLDIPQPQRQHKYVWQQYLEQFNEIKKMHFEELKTVGEISKELNISDWIILDL
FKSQNVDKLSFQELSKRRRTKDFAFLYDLHFNKKMSLKEISRAFDYSPPYIRQVFKDQGIKHLTFKNQYKN

#### Goe 14 integrase in the prophage genome

>Goe14 propjage integrase

TTGCTTATATTAAAGGGAGATCAAAAGCTAAGTACGGACAATATAATGCATTTGATTGGCAAAATTAATGCATT
GATTTATGCCAGAGTATCAACAACAGATCAAGCTAATAAGGGTTTTTCAATTGAGTCACAAATAGAAAGGTGT
AAAGAAAGAGCAATCTCTAAATTTGGATACAAGGAAAGCGAAATAATTGCGTTAGTTGAACCGGGGGGCATG
GGAGATGATCCAAACCGGCCAGCTCTTAATCACGCCTTATATTTATTAGAAAAAGGACTAGGAAAAAAATTCAT
AGTATTACACCCAGATCGTCTAACTAGAGATAACACCTTACAGGGTGTGGTATCAAGAAGAATATGGGGTATG
GGTGTTGATATCGAGTTTATTGAATTTGAAGTAAATCCTCATGATCCTGAATCAATGCTGATGTACAACATTCA
AGGTTCAATTGCACAGTACAATAAGGCGAAAATTCATGCTAACTCAAAGCGAGGAAGATTGGCTAAAGCAAA
GAAAGGTGAATTTCCTTCATTTAAAAGGTTGTATGGGTACAAATTTAATACAGTGACAGATCTTCCTGAGTATA
ACGAAGAAGAAAAAGAAATTTTGCTTGAAATGAAGGATATGCTCTTAAACAAAAAAATGTCCTCGAATGAAAT
TGCTAAAGAGCTTTCAAGAAGGGGAGTTGCTGCTCCTAACGGAAAGACTTGGTATCAGGCAACTGTAAGTAG
AATGTTGCAGAATGAAGATTACACAGGTGATTTTTATTATGGAAAGTCAAAAGTTGTTCAAATTAATGGTAAGA
AGAAACAGGTGCCGACAAACAAAGACGAATGGATATTGATAAAGATTCCTCCAATGTGGGATAAAGCTACAC
GAGAGCAAATCATTGAGCGATTGAAGAGCAACTTTAAGGGAAGAAGTCGAGCTACTAAAGACTATTTGCTGA
AAAACAAGGCGAAATGTGGTCGTTGTGGAGGGGCATGTGGCTCAGGAATAACTTCTAAAACTAAATCAGGCG
TATATAAGTATTATTCTTGTAGAGCAAAGACTGCAAAAGGATATCAAAATGGCAAGAAAGTTGTTTCATGTGA
AGGTAAGAACTGGAGAGTCGATATCGTTGATGAAGTTTTTTGGAACTGGTTTATAAAACTCCTGAAGAATCCA
GAGAAATTTCTAGAATCATTTTTAGAGGAAGCGTCTGATCAAAAGAAAATCGATGAACTCAAAGCAAAAGCAA
GTCGACTAGAAAAACAACTCGGTGAGATAGATGAAGAAATAGCAAATTATGTAATTCTTTTTGGGAAAGGGAA
AATTAAAGAAAGTATGTTCGACCAACTGTCTCAGCCATTGGAACAGAACAAAGAGCATATTGAGAATGAGATT
AAAATAATTAATTCACAATTGGCTGCAAATAAACAAACTGAAGACAAAAAACAAAAAATGATCGAGTATATTG
GCTCGTTCTCTAAAATGATTAAAAATGAAATTACAATAGAAGAAAAACGGCAATTCCTTGATTTCTTCATTGAA
AAAGTAACTCTGTTTGATGATGATCACATGGAAGTTGTATGGAAAAGTTCTTCACTTAATAATGAAAACAGCCA
TGTGGATTTTCTTAACGGCGAGCAGGGAGGGGAGTTTATAAACTCGCATAAAAGACTAAACCATATTCAAGCA
TATGGAAGACAAACAGCATGA

>Goe14 prophage integrase

LLILKGDQKLSTDNIMHLIGKINALIYARVSTTDQANKGFSIESQIERCKERAISKFGYKESEIIALVEPGGMGDDPNRP
ALNHALYLLEKGLGKKFIVLHPDRLTRDNTLQGVVSRRIWGMGVDIEFIEFEVNPHDPESMLMYNIQGSIAQYNKAK
IHANSKRGRLAKAKKGEFPSFKRLYGYKFNTVTDLPEYNEEEKEILLEMKDMLLNKKMSSNEIAKELSRRGVAAPNGK
TWYQATVSRMLQNEDYTGDFYYGKSKVVQINGKKKQVPTNKDEWILIKIPPMWDKATREQIIERLKSNFKGRSRAT
KDYLLKNKAKCGRCGGACGSGITSKTKSGVYKYYSCRAKTAKGYQNGKKVVSCEGKNWRVDIVDEVFWNWFIKLL
KNPEKFLESFLEEASDQKKIDELKAKASRLEKQLGEIDEEIANYVILFGKGKIKESMFDQLSQPLEQNKEHIENEIKIINS
QLAANKQTEDKKQKMIEYIGSFSKMIKNEITIEEKRQFLDFFIEKVTLFDDDHMEVVWKSSSLNNENSHVDFLNGEQ
GGEFINSHKRLNHIQAYGRQTA

#### A protein consisting of the C-terminal sequence from the Goe14 integrase upon prophage formation

>‘Goe14 integrase

ATGCACTTTGAAGAGCTTAAAACTGTAGGAGAGATTTCTAAGGAACTAAATATCTCAGATTGGATTATTTTAGA
TTTATTTAAATCACAAAATGTGGACAAGCTTTCATTTCAGGAATTGTCTAAGCGTAGACGAACTAAGGACTTTG
CTTTTCTTTATGATCTGCATTTCAATAAGAAAATGAGCCTGAAGGAAATTAGTCGAGCTTTCGATTATTCTCCAC
CTTATATCCGTCAAGTTTTTAAGGATCAGGGAATTAAACACTTAACCTTTAAAAATCAGTATAAAAATTAA

>‘Goe14 integrase

MHFEELKTVGEISKELNISDWIILDLFKSQNVDKLSFQELSKRRRTKDFAFLYDLHFNKKMSLKEISRAFDYSPPYIRQV FKDQGIKHLTFKNQYKN

### 8. Domain prediction in diverse SPβ integrases

#### Predicted functional domains in the integrases of SPβ, Goe14 and Lzh-a42

##### Predicted functional domains in the integrase of SPβ

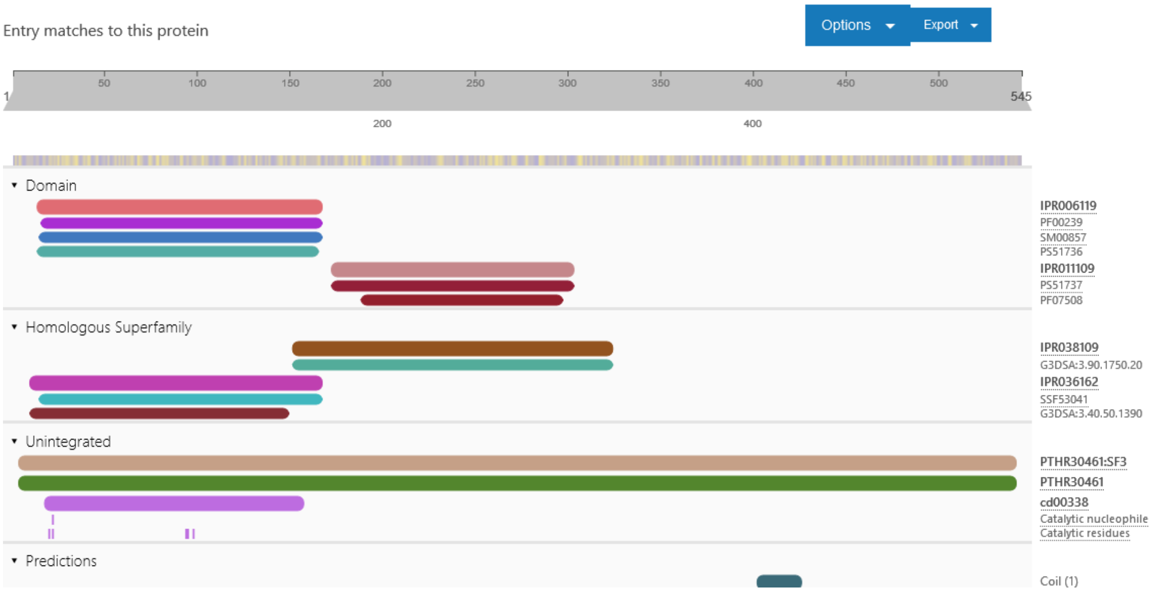

##### Predicted functional domains in the integrase of Goe14

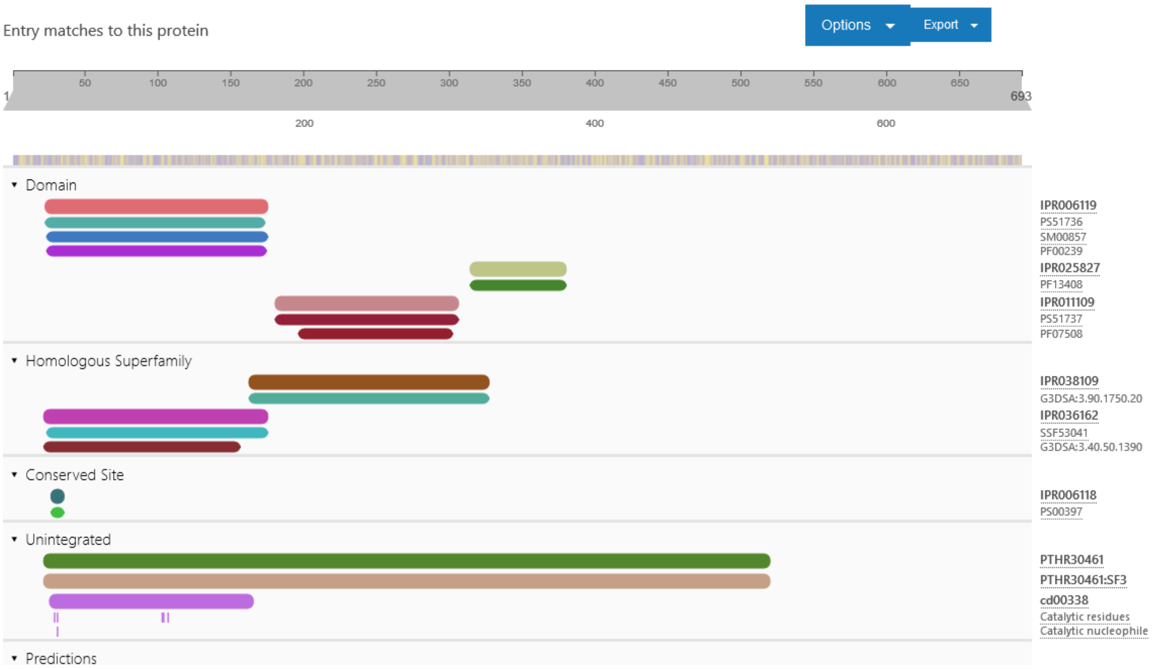

Predicted functional domains in the integrase of Lzh-a42

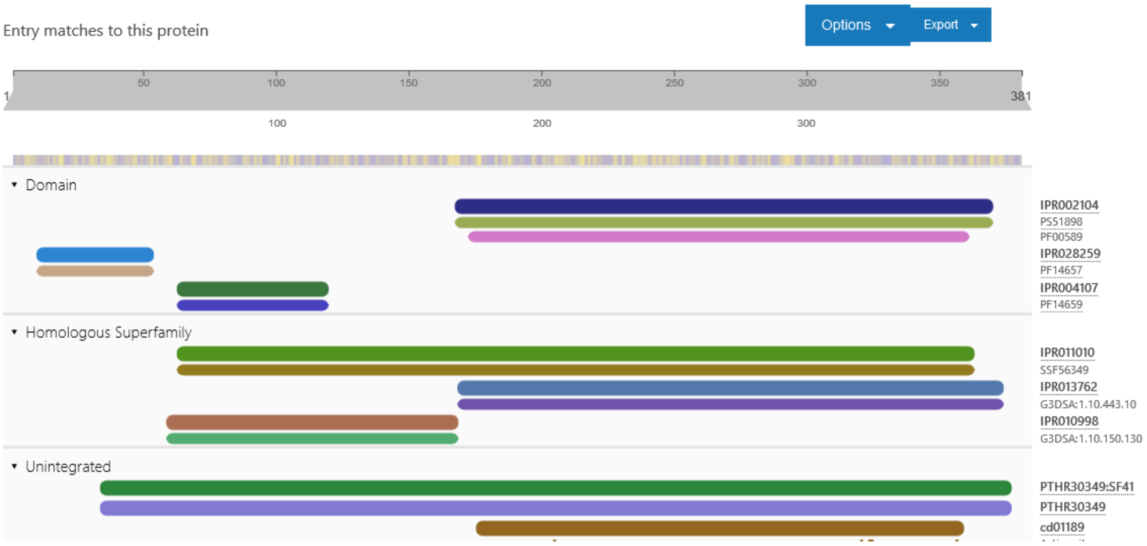

### 9. Genome integrity verification of the clear plaque mutants cpm1-6

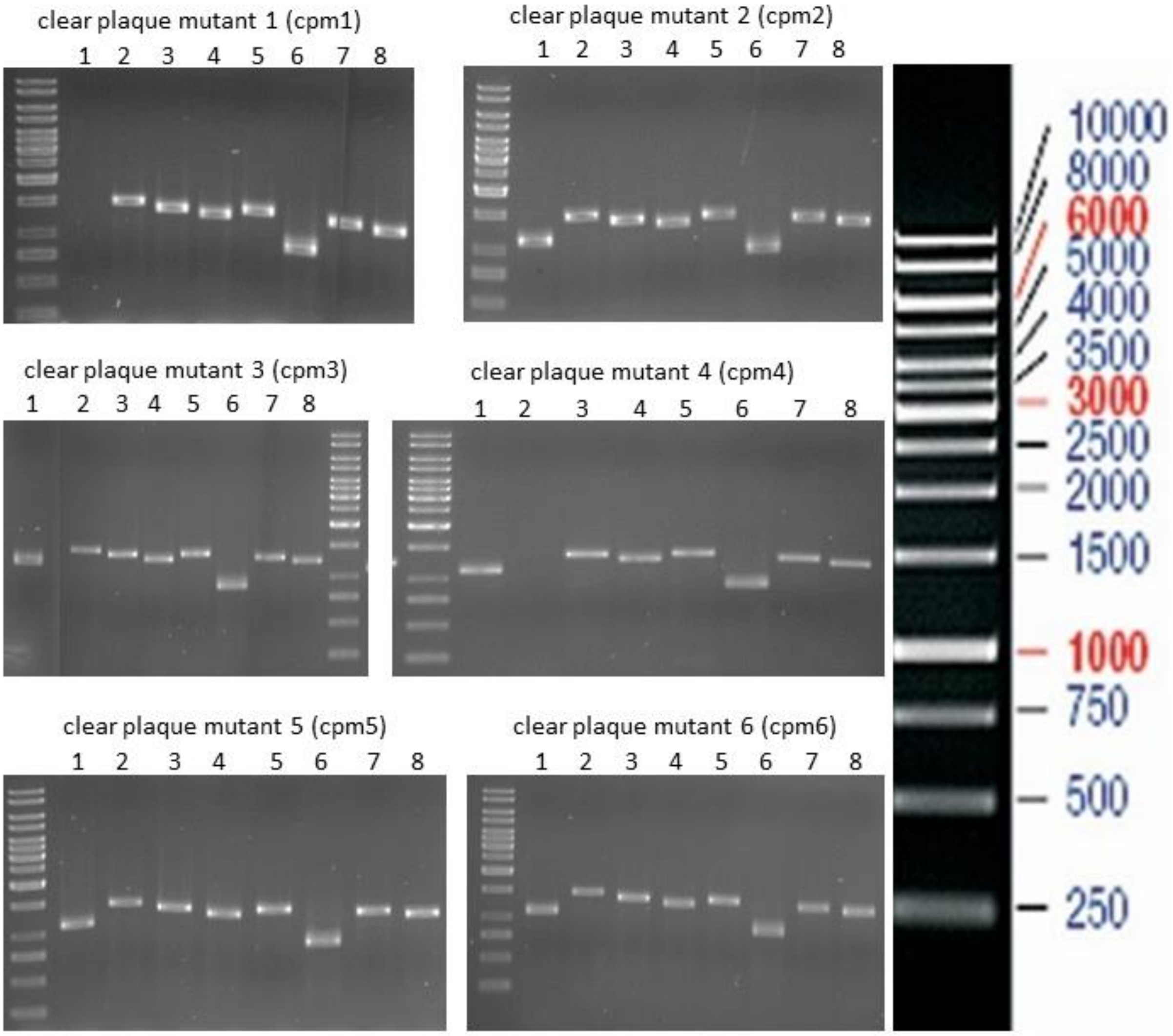

***Genome integrity verification of the clear plaque mutants cpm1-6.*** *The viral genome integrity was verified via PCR by amplifying the yorN gene with the primer PP316 / PP317 leding to a 1103 bp DNA fragment, yopR gene with the primer PP318 / PP319 leding to a 1478 bp DNA fragment, yonJ gene with the primer PP320 / PP321 leding to a 1370 bp DNA fragment, yomI gene with the primer PP322 / PP323 leding to a 1287 bp DNA fragment, yolE gene with the primer PP324 / PP325 leding to a 1383 bp DNA fragment, yokI gene with the primer PP312 / PP326 leding to a 892 bp DNA fragment, yokE gene with the primer PP327 / PP328 leding to a 1303 bp DNA fragment, and yosL gene with the primer PP359 / PP360 leding to a 1500 bp DNA fragment.*

### 10. Clear plaque mutant compmelentation

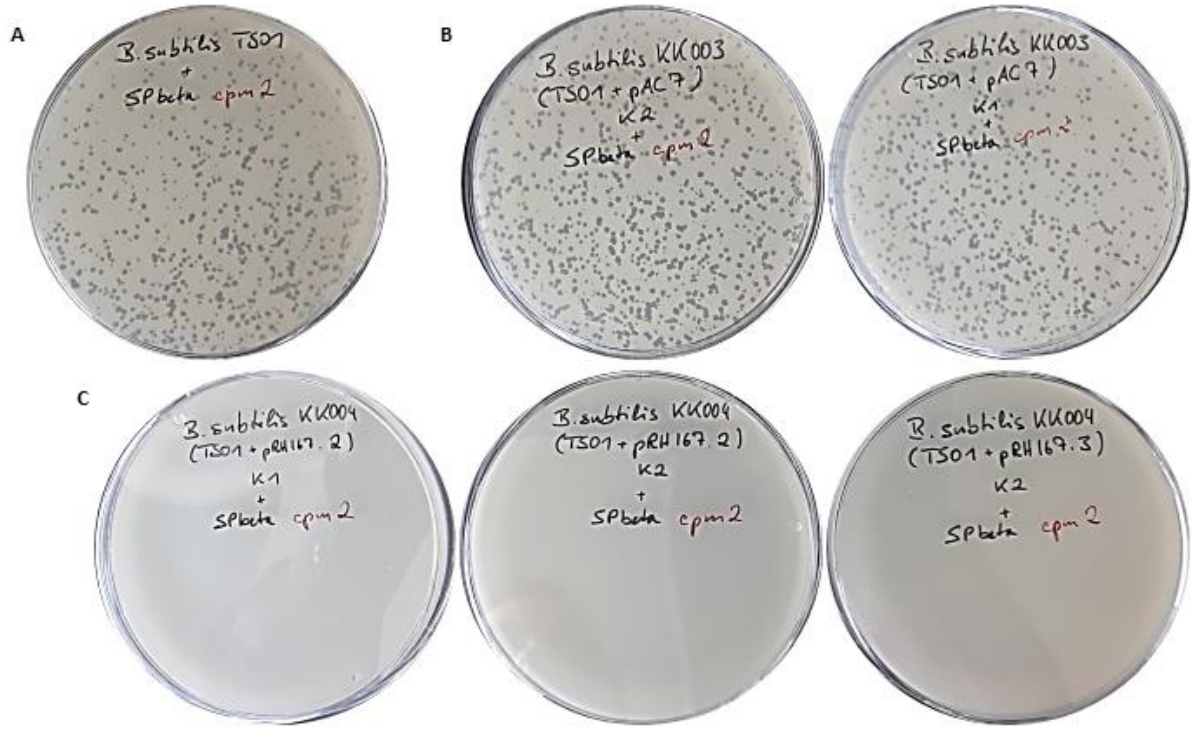

**Plaque assay with the clear plaque mutant SPβ cpm2.** The used amount of phages was always the same for the infection with each host strain. **A:** Controllexperiment with *B. subtilis* TS01. SPβ cpm2 lytically replicates and forms plaques. **B:** Controllexperiment with *B. subtilis* KK003, which was transformed with the empty vector pAC7. SPβ cpm2 lytically replicates and forms plaques with a comparable efficiency like with *B. subtilis* TS01. **C:** Complementation experiments with *B. subtilis* KK004 clones, which artificially expresses YopR. The SPβ cpm2 phage cannot form plaques on this host.

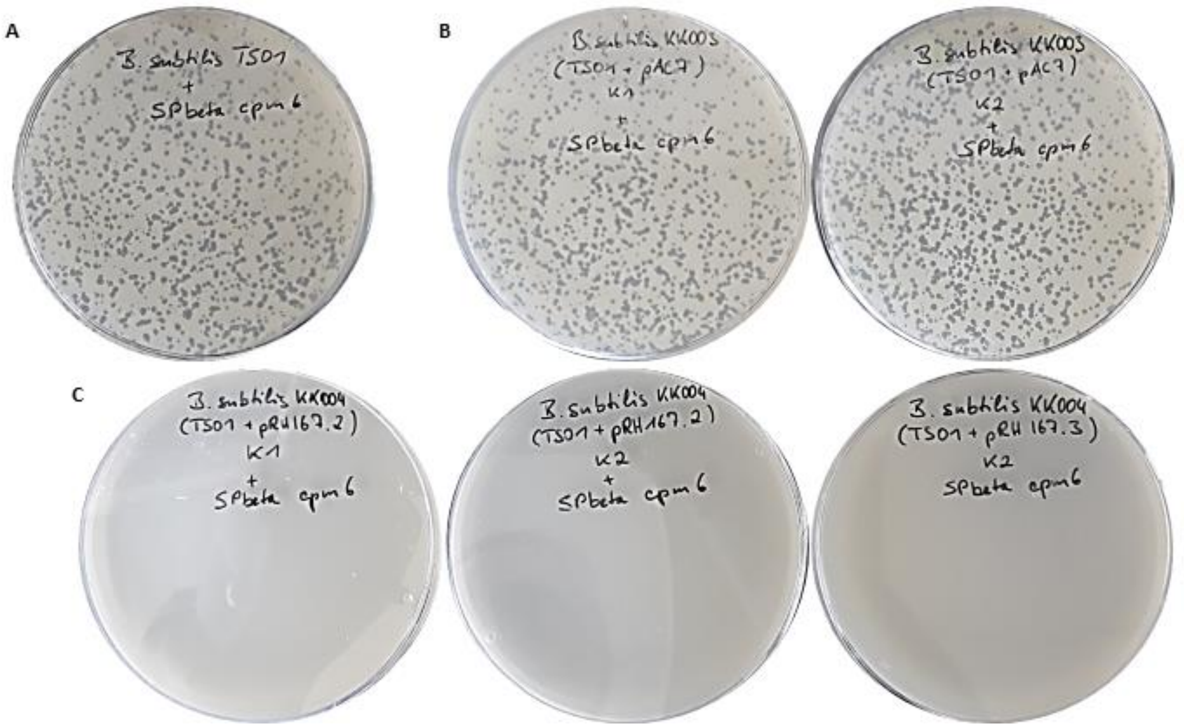

**Plaque assay with the clear plaque mutant SPβ cpm6.** The used amount of phages was always the same for the infection with each host strain. **A:** Controllexperiment with *B. subtilis* TS01. SPβ cpm6 lytically replicates and forms plaques. **B:** Controllexperiment with *B. subtilis* KK003, which was transformed with the empty vector pAC7. SPβ cpm2 lytically replicates and forms plaques with a comparable efficiency like with *B. subtilis* TS01. **C:** Complementation experiments with *B. subtilis* KK004 clones, which artificially expresses YopR. The SPβ cpm2 phage cannot form plaques on this host.

### 11. A ~250bp genome fragment containing the SPβ-like pac-site

#### Genome fragment containing the SPβ pac-site (<-)

GCTATGGGTACATCTGTTCCCTTAATTTTACCATTAGATGTAAATATACCCCCATATATTGGTATTGTAATGCGT
GTATAAGGAACATTATACGTGCATTTAATGACTTGGGCAGGTCTGCACCAATGCTATCATAATTCCAGGTCAAA
CACAAGAATTAATTTTTAAAAAATAAATTGTTTTTCAATTTAAAAATGAATAACCATTCATTAAAAATAATCGTA
TTTATAACGAATCCGAGGGAATCGAAG

#### Genome fragment containing the phi3T pac-site (<-)

GCTATGGGTACATTCGTTCCCTTAATTTCACCATTAGATGTTAATATACCCCCTGTATATTGGTATTGAATTCCCT
ATATAAGATACGTTATGTAGGGTTTTTATAATTTGGCTGATAAATTTGCACAGCTAACACAATTTCAAATGAAA
AACAATAGTTATTTTTGAAAAAATAATTAAGTGGTCAATTTAAAAGTGAATACCCATTCATTAAAAATAATCGT
GTATATAACGAAAGGAAGCAAATCGAAG

#### Genome fragment containing the Goe11 pac-site (<-)

GCTATGGGTACATTTGTTCCCTTTATTTTACCATTAGATGTAAATATACCCCCATATATTGGTATTGTAATACGTG
TATAAGGAACATTATACGTGTATTTGATGACTTGGGTAGGCCTACAACAGTGCTAATATAATTCCAGGCCAAAC
ACAAGAATTAATTTTAAAAAAAATAAATTGGTTATCAATTTAAAAATGAATAACCATTCATTAAAAATAATCGTA
TTTATAACGAATCCGAGGGAATCGAAG

#### Genome fragment containing the Goe14 pac-site (<-)

GCTATGGGTACATTTGTTCCCTTTATTTTACCATTAGATGTAAATATACCCCCATATATTGGTATTGTAATACGTG
TATAAGGAACATTATACGTGTATTTGATGACTTGGGTAGGCCTACAACAGTGCTAATATAATTCCAGGCCAAAC
ACAAGAATTAATTTTAAAAAAAATAAATTGGTTATCAATTTAAAAATGAATAACCATTCATCAAAAATAATCGT
ATTTATAACGAATCCGAGGGAATCGAAG

#### Genome fragment containing the Goe12 pac-site (<-)

GCTATGGGTACATCCGTTCCCTTTATTTTACCATTAGATGTAAATATACCCCCATATATTGGTATTGTAATACGT
GTATAAGGAACATTATACGTGCATTTAATGGTTTAGATATGTTTACAACAGTGCTAACATAATTCCAGGCCAAA
CACAAGAATAAATTTTTAAAAAAATAAATTGGTTATCGATTTGAAAAGAATAACCATTCATCAAAAATAATCGT
ATTTATAACGAATCCGAGGGAATCGAAG

#### Genome fragment containing the Goe13 pac-site (<-)

GCTATGGGTACATCCGTTCCCTTTATTTTACCATTAGATGTAAATATACCCCCATATATTGGTATTGTAATACGT
GTATAAGGAACATTATACGTGCATTTAATGGTTTAGATATGTTTACAACAGTGCTAACATAATTCCAGGCCAAA
CACAAGAATAAATTTTTAAAAAAATAAATTGGTTATCGATTTGAAAAGAATAACCATTCATCAAAAATAATCGT
ATTTATAACGAATCCGAGGGAATCGAAG

#### Multiple sequence alignment all verified genome fragment containing the SPβ pac-site

```
CLUSTAL O(1.2.4) multiple sequence alignment

phi3T           GCTATGGGTACATTCGTTCCCTTAATTTCACCATTAGATGTTAATATACCCCCTGTATAT60
SPβ             GCTATGGGTACATCTGTTCCCTTAATTTTACCATTAGATGTAAATATACCCCCA-TATAT59
Goe11           GCTATGGGTACATTTGTTCCCTTTATTTTACCATTAGATGTAAATATACCCCCA-TATAT59
Goe14           GCTATGGGTACATTTGTTCCCTTTATTTTACCATTAGATGTAAATATACCCCCA-TATAT59
                *************  ******** **** ************ ***********  *****

phi3T           TGGTATTGAATTCCCTATATAAGATACGTTATGTAGGGTTTTTATAATTTGGCTGATAAA120
SPβ             TGGTATTGTAATGCGTGTATAAGGAACATTATACGTGCATTTAATGACTTGGGCAGGTCT119
Goe11           TGGTATTGTAATACGTGTATAAGGAACATTATACGTGTATTTGATGACTTGGGTAGGCCT119
Goe14           TGGTATTGTAATACGTGTATAAGGAACATTATACGTGTATTTGATGACTTGGGTAGGCCT119
                ******** * * * * ******  ** ****    *  *** ** * ****
```

```
phi3T       TTTGCACAGCTAACACAATTTCAAATGAAAAACAATAGTT-ATTTTTGAAAAAATAATTA179
SPβ         GCACCAATGCTATCATAATTCCAGGTCAAACACAAGAATTAATTTTTAA-AAAATAAATT178
Goe11       ACAACAGTGCTAATATAATTCCAGGCCAAACACAAGAATTAATTTTAAAAAAAATAAATT179
Goe14       ACAACAGTGCTAATATAATTCCAGGCCAAACACAAGAATTAATTTTAAAAAAAATAAATT179
                **  ****  * **** **    *** **** * ** *****  * ******* *

phi3T       AGTGGTCAATTTAAAAGTGAATACCCATTCATTAAAAATAATCGTGTATATAACGAAAGG239
SPβ         GTTTTTCAATTTAAAAATGAATAACCATTCATTAAAAATAATCGTATTTATAACGAATCC238
Goe11       GGTTATCAATTTAAAAATGAATAACCATTCATTAAAAATAATCGTATTTATAACGAATCC239
Goe14       GGTTATCAATTTAAAAATGAATAACCATTCATCAAAAATAATCGTATTTATAACGAATCC239
              *  *********** ****** ******** ************ * *********

phi3T       AAGCAAATCGAAG 252
SPβ         GAGGGAATCGAAG 251
Goe11       GAGGGAATCGAAG 252
Goe14       GAGGGAATCGAAG 252
             **  ********
```

#### SPβ pac-site region

```
GCTATGGGTACATCTGTTCCCTTAATTTTACCATTAGATGTAAATATACCCCCATATATTGGTATTGTAATGCGT
.....(((.((....)).)))((((((((((((.((.(((............))).)).)))))(((((((((((

GTATAAGGAACATTATACGTGCATTTAATGACTTGGGCAGGTCTGCACCAATGCTATCATAATTCCAGGTCAAAC
((((((......)))))))))))))...((((((((((((((....)))..)))..........))))))))...

ACAAGAATTAATTTTTAAAAAATAAATTGTTTTTCAATTTAAAAATGAATAACCATTCATTAAAAATAATCGTAT
)))))))))))...........(((((((.....)))))))..(((((((....)))))))..(((((....)))

TTATAACGAATCCGAGGGAATCGAAG
))....(((.(((...))).)))...
```

Inverted repeats in the ~250 bp genome fragment containing the SPβ pac-site

### Visualisation of the predicted inverted repeats in the predicted SPβ pac-site region

The sequence is presented in 5’>3’ orientation. The packaging direction is counterclockwise, meaning the start point is at position 251.

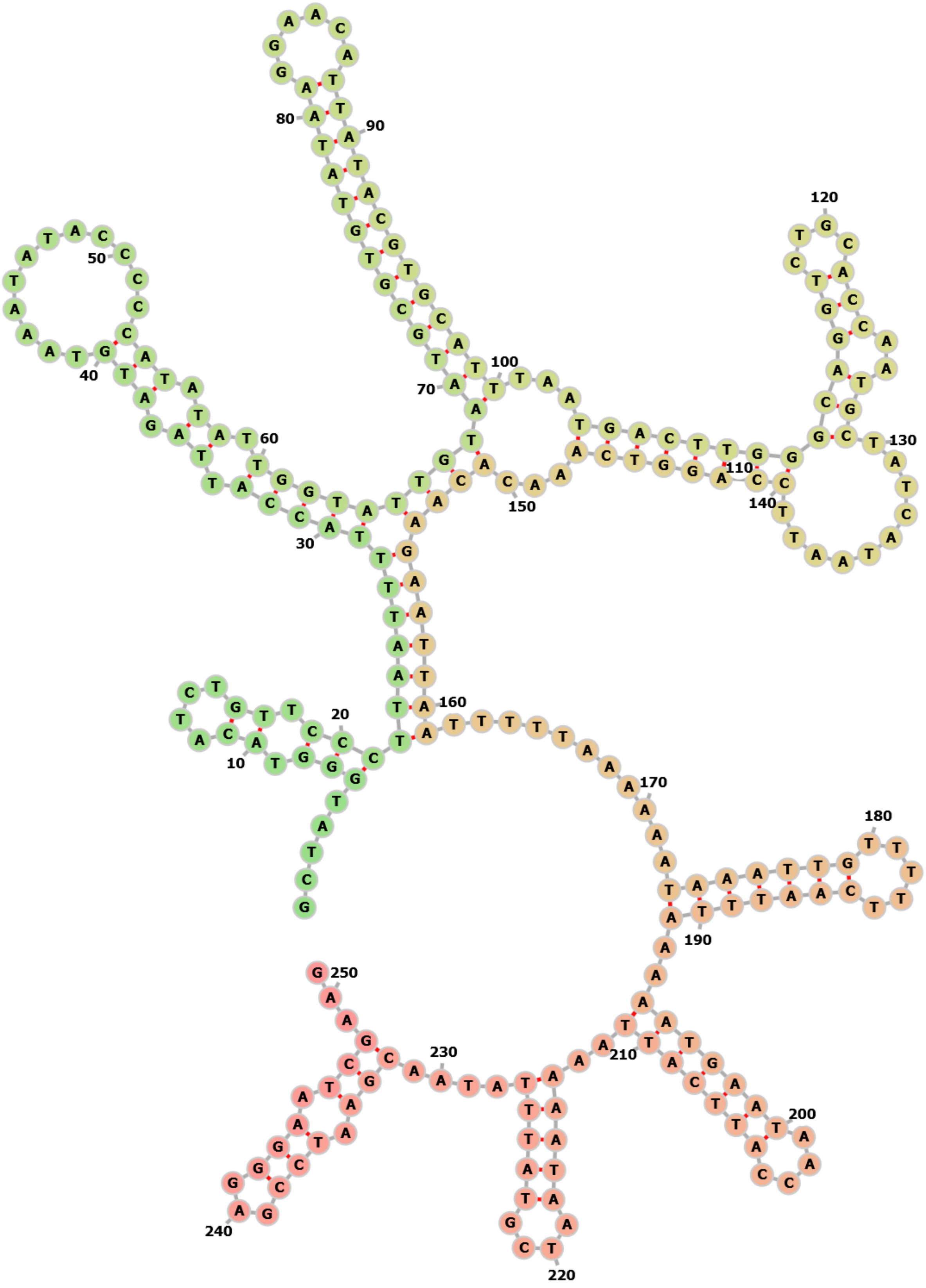
